# Mesoscale Functional Reorganization of Cortical Networks After Cortical Spreading Depression

**DOI:** 10.64898/2026.05.24.727472

**Authors:** Bengisu Solgun, Aslıhan Bahadır-Varol, Buket Dönmez-Demir, Engin Demir, Hülya Karataş, Şefik Evren Erdener

## Abstract

**Background:** Resting-state functional imaging is increasingly used to understand how cortical networks modulate and respond to pain. Awake imaging with a minimally invasive approach is key to observe the natural state of the brain. As migraine with aura, a common headache disorder, can be experimentally modeled by cortical spreading depressions (CSD) in rodents, it is essential to understand the impact of CSD on functional connectivity and network topology to find imaging cues of trigeminovascular activation and headache.

**Methods:** We used awake widefield intrinsic optical-signal imaging (IOSI) on optically cleared windows to non-invasively characterize the impact of CSDs on bihemispheric resting-state static and dynamic functional connectivity patterns and network topology. A subset of mice was chronically treated with amitriptyline to examine the effect of susceptibility to CSD on connectivity. After baseline imaging, CSD was triggered optogenetically and confirmed by laser speckle contrast imaging. A group of mice received intraperitoneal naproxen after CSD to suppress headache. IOSI was repeated at 30 minutes, 60 minutes, 4 hours, and 24 hours after CSD. The mouse grimace scale was scored at each time point for behavioral headache documentation.

**Results:** We observed time-dependent changes in resting-state functional connectivity that were reversed by naproxen. Amitriptyline, a prophylactic migraine medication, decreased susceptibility to CSD and modified resting-state functional connectivity differently than controls. Network analysis with graph-theoretical methods revealed barrel and retrosplenial cortices as potential key players in trigeminal pain processing after CSD. Dynamic functional connectivity analysis demonstrated functional connectivity states, with fractional occupancy and mean dwell time of these states showing distinct CSD and pain-modulated states. A support vector machine was utilized to predict CSD-mediated dynamic connectivity changes in controls.

**Conclusion:** Our results bring insight into potentially headache-associated changes in resting-state cortical functional connectivity after CSD and how this functional reorganization is influenced by acute and chronic medications for migraine.

## Introduction

Migraine is a common headache disorder characterized by a unilateral throbbing headache accompanied by nausea, photophobia, phonophobia, and is preceded by aura in approximately 20% of cases ^1–3^. Although migraine has a significant social, clinical, and economic impact, presently, diagnosis relies on clinical criteria ^1,4–7^. An increasing number of studies are conducted in search of a reliable marker for diagnosis and monitoring of treatment response ^8^. Among modalities, resting-state functional imaging is promising to gain a better understanding of cortical pain processing. While prior evidence from functional imaging in migraine patients suggests modulation of various cortical areas in response to trigeminovascular activation, reported findings are inconsistent across studies and the impact of headache pain on large scale cortical functional organization remains poorly understood ^9–16^. Nevertheless, each study introduces multiple regions as potential key players, reintroducing migraine as a network disease rather than an isolated local dysfunction in the brain ^9,10^.

To address this knowledge gap, migraine aura can be modeled by cortical spreading depression (CSD) in rodents ^17^. CSD, the underlying mechanism of migraine aura, is a slow self-propagating wave of depression followed by depression of cortical activity, which by nature disrupts cerebral hemodynamics substantially and alters blood flow through neurovascular coupling ^17–20^. The effect of CSD on cerebral hemodynamics and functional connectivity immediately following CSD has been studied ^21–23^, but the impact of the resulting headache and its trigeminal processing on cortical functional networks is currently underexplored. In addition, prior resting functional connectivity studies in rodents primarily relied on anesthesia and invasive approaches for CSD induction and imaging ^23,24^ and research involving awake animals and minimally invasive cranial windows have been limited.

Prophylactic migraine medications such as amitriptyline, propranolol, or valproate commonly used by patients have been shown to decrease susceptibility to CSD in rodents ^25^. While this can be a mechanism for their therapeutic effect in decreasing migraine attack frequency, it is also possible that these medications could be affecting the functional connectivity parameters and network topology, either in baseline conditions or in response to CSD, relieving migraine headaches. Investigation of these interactions could help better characterization of migraine-specific effects of these prophylactic medications with complex pharmacodynamics.

In this study, we investigate the impact of headache induction following CSD on resting-state functional connectivity and network topology in awake mice using intrinsic optical signal imaging (IOSI). We show that CSD leads to distinct alterations in the cortical functional architecture at resting-state, accompanied by headache-indicating mouse grimace scale scores and that these alterations are reversed by naproxen, an analgesic medication. We also explore how amitriptyline, a prophylactic medication for migraine, modulates this network response to CSD. In parallel, we investigate if susceptibility to CSD is reflected on baseline functional connectivity. Lastly, we implement a dynamic connectivity approach to reveal CSD and pain modulated states and we utilize a support vector machine to classify dynamic functional connectivity values before and after CSD. Our findings demonstrate distinct alterations to the resting-state cortical functional network with CSD, and with medications for migraine.

## Materials and Methods

### Animals

2-4-month-old C57BL/6J-Thy1-COP4/EYFP mice (n=52) (The Jackson Laboratory, strain: 007612) expressing light-activated Channelrhodopsin-2 in excitatory neurons were used. 2 mice were excluded from the study: one due to CSD being triggered twice and one due to incorrect headplate placement. Mice were housed at stable temperature (19-22 °C), 12:12 light/dark cycle with ad libitum access to food and water. All experiments were approved by the Hacettepe University Animal Experimentations Local Ethics Board (Approval number: 2022/40).

Prior to experiments, mice were randomly separated into chronic amitriptyline-administered, naproxen-administered and control groups. Experimental timeline can be seen in Figure 1A.

**Fig. 1.**
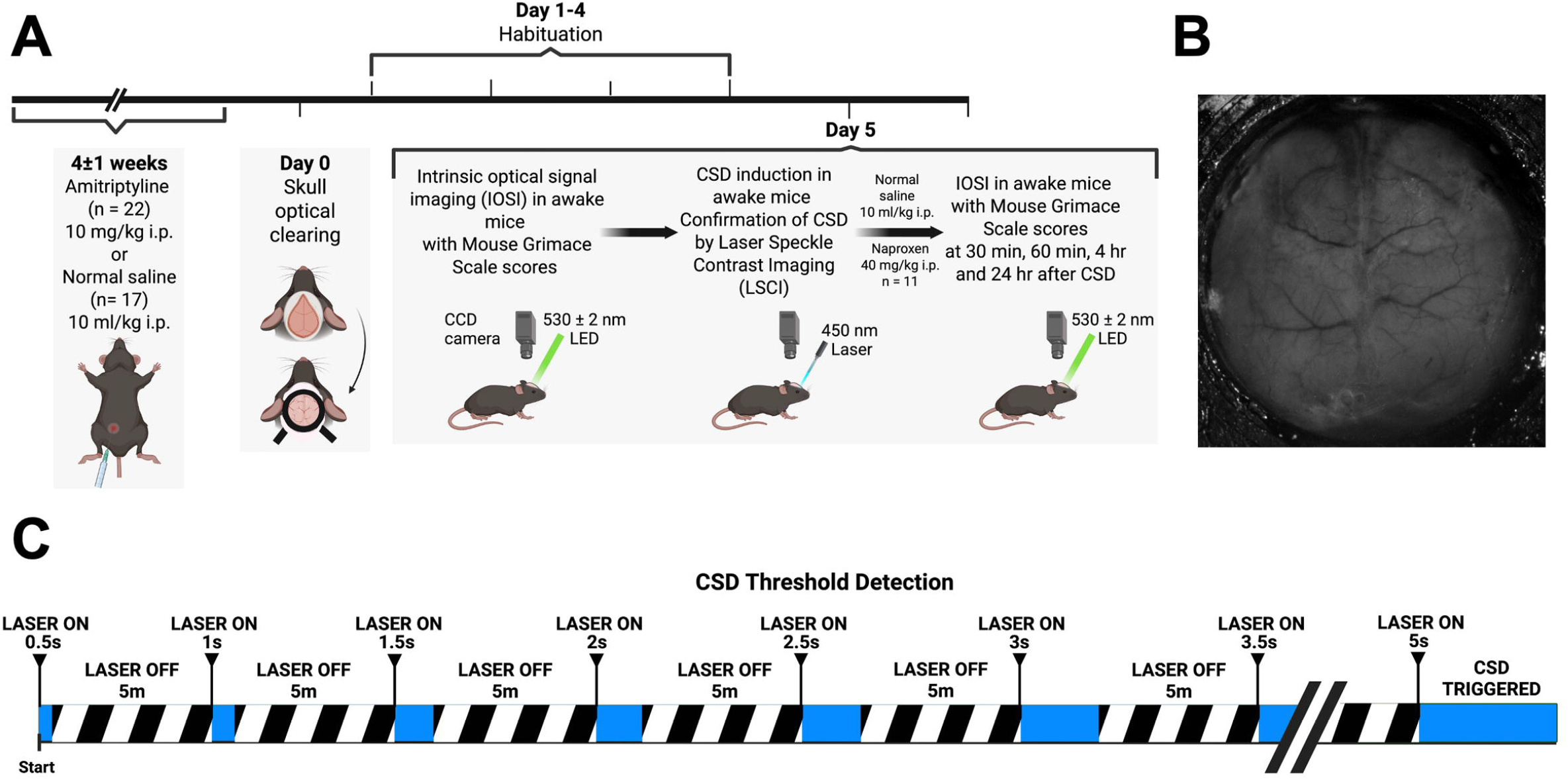
**(A)** Experimental timeline **(B)** Raw IOS image of an optically cleared skull. **(C)** CSD threshold was detected by increasing the duration by 0.5 seconds every 5 minutes until a CSD was triggered.

### Surgery and skull optical clearing

Mice were anesthetized with isoflurane (4% induction, 1-2% maintenance in oxygen), head-fixed in a stereotaxic frame (World Precision Instruments, USA) under a stereomicroscope (SMZ745T, Nikon Instruments Inc., Japan). Temperature was monitored with a rectal probe and maintained at 37.0°C on a homeothermic blanket control unit (Kent Scientific, USA). Eye ointment (Recugel, Bausch + Lomb, Canada) was applied throughout surgery. Hindlimb reflexes were periodically checked to evaluate the depth of anesthesia. We implemented a minimally invasive, ready-to-image, chronic transparent skull technique suitable for awake mouse optical imaging, as described elsewhere ^26^. Following hair removal and disinfection of the scalp with 70% ethanol and 10% povidone-iodine, a midline incision was made to the scalp, large enough to visualize both hemispheres. Both temporalis muscles were retracted from their insertion, avoiding damage to the superficial temporal vein ^27^ and the remaining groove was filled with cyanoacrylate glue (Loctite 401, Henkel, Germany) to provide a solid surface for head plate adhesion. The skull was dried with a clean pressurized air spray before head plate adhesion. Next, the skull was cleared by the topical application of 10% EDTA in 1X phosphate-buffered saline (PBS) solution, followed by the application of NOA61, a UV-curable adhesive ^26,28–30^. The transparent window was then covered with a custom-cut coverslip to fit the final window size (10 mm diameter). NOA61-applied window was cured under 310-330 nm UV light to obtain a solid, stable transparent window. After clearing, the window was immediately closed with a custom-made cap to prevent light exposure to Channelrhodopsin-2 positive transgenic animals. Cyanoacrylate and dental cement were applied to cover all exposed skull around the window. After surgery, the animal was returned to a clean cage, placed on a heating pad for recovery, and monitored until fully recovered from anesthesia. Ibuprofen in drinking water (50mg/100ml) was provided for post-operative analgesia for 1 day. A sample transparent window ready for imaging at 5 days after skull optical clearing can be seen in Figure 1B.

### Habituation to the Imaging Setup for Awake Head-Fixed Imaging

Habituation to the setup for awake imaging was started 1 day after surgery. Four habituation sessions (1 session/day), with durations of 10, 20, 30, and 40 minutes, were performed, with the mouse head fixed in the imaging cradle and rewarded every 5 minutes with sweetened condensed milk. If the mouse showed any signs of restlessness or visible stress during habituation, the session was terminated, and the mouse was returned to its home cage.

### Induction of CSDs

A 450nm laser coupled to a fiberoptic cable (diameter: 200μm, numerical aperture: 0.53, optical power at illumination spot: 6 mW, Doric Lenses, Canada) was placed on the transparent window in the right frontal area at approximately a 60**°** angle to the skull. Starting with a 0.5 s stimulus, stimulus duration was increased every 5 minutes until a CSD wave was observed and confirmed with laser speckle contrast imaging (LSCI) (Figure 1C, 3A, 3B) ^18,31^.

### Drug Administration

Amitriptyline-administered group received i.p. amitriptyline (10 mg/kg, 1 mg/ml solution in saline), while the remaining groups received normal saline (10 ml/kg) daily for 4±1 weeks prior to skull optical clearing, continued until final imaging session ^25^.

Naproxen-sodium (40 mg/kg ^32^, Sigma-Aldrich, USA), a prostaglandin synthesis inhibitor, was administered intraperitoneally (4 mg/ml solution in saline) to the naproxen-administered group, 3 to 5 minutes after CSD onset, to modulate headache. The remaining groups (amitriptyline-administered and control groups) received normal saline at this time.

### Imaging

All imaging sessions were performed with the mice fully awake, head-fixed in the imaging cradle; brief anesthesia (4% isoflurane) was only used to place the animals in the imaging cradle. A stereomicroscope (SMZ1000, Nikon Instruments Inc., Japan) mounted with a CCD camera (Basler acA1300-60gmNIR, Basler Vision Technologies, Germany) was used for IOSI and LSCI. We used a 530±2 nm LED light (Thorlabs, USA) for IOSI. 530nm is near the isosbestic point for hemoglobin, and IOS imaging at a single wavelength for total hemoglobin produces comparable resting-state functional connectivity maps to 3 wavelength imaging for oxy-, deoxy-, and total hemoglobin ^33^. 8-bit 512×512-pixel images were obtained through 2×2 binning of 1024×1024-pixel images. For laser speckle contrast imaging, the window was illuminated using a 785-nm laser diode (Thorlabs, USA); 8-bit 1276×1020-pixel images were obtained.

### Image Processing

For LSCI, image sets (15 raw speckle images, with an exposure time of 5 ms) were averaged. Using a sliding window of 7 × 7 pixels, a spatial speckle contrast (K) image was computed. Speckle contrast images were converted to integrated correlation time (ICT) images (1/K^2^) to obtain blood flow index values. An open-access MATLAB script was used to process IOS images ^33^. 512×512 images were downsampled to 128×128 pixels, temporally detrended, brain-masked, bandpass-filtered (0.035-0.08Hz), and globally regressed.

### Static functional connectivity analysis

ROI coordinates were determined according to Allen Brain Atlas with respect to manually selected bregma coordinates ^34–36^. 10 regions of interest (5 in each hemisphere) were selected for analysis: secondary motor areas, primary sensory areas (barrel field, hindlimb, forelimb) and retrosplenial areas based on their potential roles in pain processing and modulation ^11,13,37–50^. For each ROI, 5 seeds were selected as follows: the atlas-determined coordinate was selected as the center seed (Seed 1), around which a 3-by-3 pixel square was created. The remaining 4 seeds were the corners of this square (Supplementary Figure 1A). For each seed, seed size was 3x3, and another 3-by-3 pixel square was created, and the values from 9 pixels in this region were averaged for each seed signal (Supplementary Figure 1A). For functional connectivity analyses between regions, all ROI seed signals were Pearson-correlated with each other, and the results were Fisher-z transformed. Next, the z values of 5 seeds corresponding to the same ROI were averaged. For functional connectivity analyses within a region, a 5-by-5 square with Seed 1 as its center was created, and every pixel in this square (25 pixels in total) was Pearson-correlated with each other. Next, the values were Fisher z-transformed and then averaged to obtain a within-region connectivity value for each ROI.

### Behavioral documentation of headache after CSD

We used the Mouse Grimace Scale (MGS), a well-established method to document facial responses in mice with pain ^51^. Due to head plate positioning, ear position, and nose bulge parameters were determined to be unreliable for scoring; MGS were scored with the remaining three parameters: orbital tightening, cheek bulge, and whisker change ^52,53^. This modified version of the MGS is hereinafter referred to as the modified MGS. Facial images were taken immediately after each 8-minute imaging session. All images were scored by 2 blinded scorers, and the scores were averaged.

### Correlation of modified MGS scores with functional connectivity

We used repeated measures correlation to showcase whether there is a reliable association between functional connectivity values and modified MGS scores for each animal ^54^. Fisher z-transformed baseline, 30-minute, 60-minute, 4-hour, and 24-hour values were included. Functional connectivity values from the left and right barrel cortex were correlated separately with modified MGS scores for each of the groups using repeated measures correlation (Pingouin v0.5.5 ^55^ in Python v3.11.8).

### Network Analysis

Anti-correlations may also contain valuable information; therefore, we chose not to discard these connections and used the absolute value of each Fisher z-transformed Pearson correlation matrix ^56–60^. We then used a minimum spanning tree to construct the skeleton of the graph with the minimum number of edges that connect all nodes ^61,62^. This approach ensures that the graphs are not fragmented, as disconnected graphs may lead to unreliable results in centrality measures ^59,63,64^. With N as the number of nodes in a graph, our minimum spanning tree had 10 nodes and 0.2 (2/N) density. Next, we used fixed-density thresholding to add more edges to the network until our goal density (0.35) was achieved (35% of the strongest edges out of all possible edges between node pairs were taken as edges), and then binarized matrices to obtain a graph for each imaging session of an animal. All network analyses were done in NetworkX v3.1 ^65^ (Python v3.11.8).

### Dynamic Functional Connectivity

To avoid spurious fluctuations in dynamic functional connectivity analysis, a window length larger than the largest wavelength (approximately 28.6 seconds in our study after bandpass filtering) is recommended, and a window length of 30-60 seconds is generally used ^66^. We used a sliding window approach to split the time series into rectangular windows (window length: 120 frames (40s), window step size: 2 frames), resulting in 661 windows per imaging session. Correlation matrices between ROI signals were computed separately for each window, and our approach was identical to static functional connectivity analyses: 9 pixels were averaged for each seed, and 5 seeds were selected for each ROI. Then, Pearson correlation values of 50 seed signals from 10 regions were computed and Fisher z-transformed. Finally, z-values of seeds corresponding to the same ROI were averaged to obtain a 10x10 functional connectivity matrix for each window per imaging session for each animal.

### Support Vector Machine Classifier

Dynamic functional connectivity values at baseline and 60 minutes were used as connectivity changes generally peaked at 60 minutes and then started returning to baseline. Functional connectivity parameters that were significantly different between baseline and 60 minutes in the control group were selected as features and normalized per animal to reduce animal-specific variations. Control mice were divided into a training (80%) set and an independent hold-out test (20%) set (functional connectivity data from 13 and 4 control animals, respectively). Within the training set, hyperparameter tuning, including feature selection (using ANOVA F-values), was conducted using leave-one-group-out cross-validation (with different animals as “groups”). 4 features were selected: Left barrel within region connectivity, left barrel-right barrel, left barrel-right retrosplenial, and left secondary motor-left retrosplenial functional connectivity (Figure 9A). The final model (an SVM with a linear kernel) optimized to maximize balanced accuracy was trained on the full training set with the optimal hyperparameters (C = 0.01) and was then evaluated on the hold-out test set of controls. The model was also used for cross-prediction on naproxen and chronic amitriptyline groups. Data labels have been randomly shuffled to train the classifier and compare model performance on the randomized dataset to the original dataset. Model performance was evaluated using AUC, accuracy, and F1 scores. SVM was trained in Scikit-learn 1.7.1 ^67^ (Python 3.11.8). The analysis pipeline can be seen in Figure 2.

**Fig. 2.**
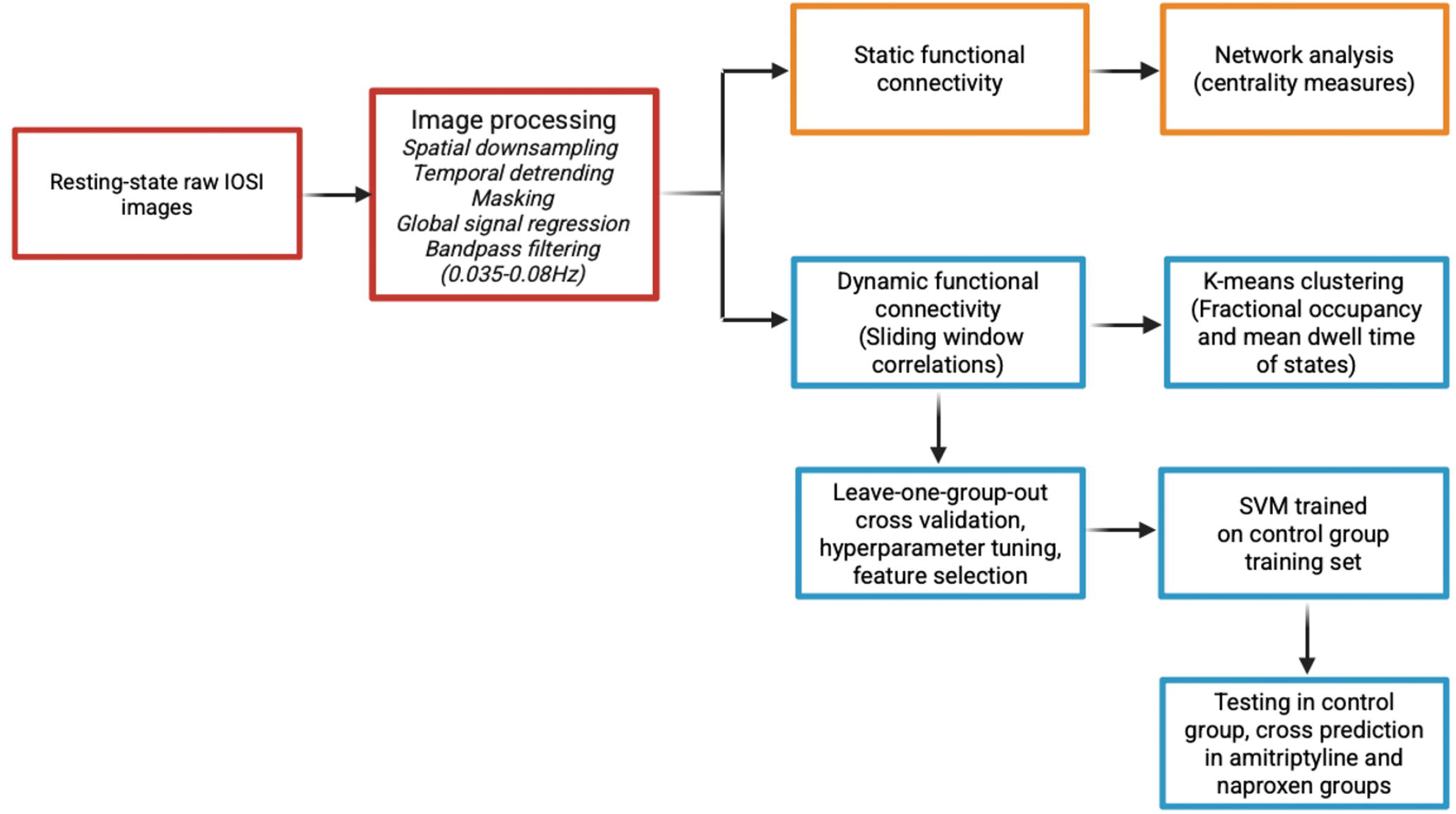
Data analysis pipeline. Image processing steps are shown in red, static functional connectivity-based analysis steps are shown in orange, and dynamic functional connectivity-based analysis steps are displayed in blue. IOSI: intrinsic optical signal imaging, SVM: support vector machine.

### K-means Clustering

Using the features that were significantly different 60 minutes after CSD compared to baseline in the control group, the K-means clustering algorithm was used to cluster dynamic functional connectivity values into states ^68^. The elbow criterion of the cluster validity index (within-cluster distance to between-cluster distance) was used to determine the optimal number of clusters (k) as 4 (Supplementary Figure 9B); the elbow point was confirmed with KneeLocator in Python. Subject exemplar windows were chosen for each imaging session of each animal based on local maxima in functional connectivity variance, resulting in 33.23± 3.63 windows per imaging session. The K-means clustering algorithm was repeated 500 times with random initialization of centroids using the subject exemplar windows to escape local minima; the resulting centroids were then used to cluster all data ^68^. Fractional occupancy and mean dwell time were computed for each state and each imaging session. Fractional occupancy is the count of how many windows each state occupies for each imaging session of an animal. These were divided by the total number of all windows of an imaging session and reported as percentages. Mean dwell time is the number of consecutive windows an animal remained in a state at each imaging session (plotted as seconds) and was also calculated for each animal at each imaging session separately.

### Statistical Analyses

Statistical analyses were performed using Graphpad Prism Version 10.3.1 (GraphPad Software, Boston, Massachusetts, USA, www.graphpad.com). All data were plotted as means ± SD. Normality of distribution was determined by the Shapiro-Wilk test. Static functional connectivity changes between regions of interest across time, static functional connectivity changes within regions of interest across time, changes in each centrality measure (degree, eigenvector and PageRank centrality) for each group (control, naproxen, chronic amitriptyline) were compared using one-way ANOVA for repeated measures (Tukey test for multiple comparisons) or Friedman test (Dunn test for multiple comparisons) and adjusted for multiple comparisons with the Benjamini-Hochberg false discovery rate (FDR) (FDR-corrected p<0.05) ^69,70^. CSD threshold values between groups were compared using a two-tailed t-test. Baseline static functional connectivity values between regions of interest, within regions of interest, and baseline values for each centrality measure between high and low threshold groups and between control/naproxen and amitriptyline groups were compared using a two-tailed t-test or Mann-Whitney U test and adjusted for multiple comparisons with the Benjamini-Hochberg false discovery rate (FDR-corrected p<0.05) ^69,70^. Mouse grimace scale scores were compared using one-way ANOVA for repeated measures (Tukey test for multiple comparisons) or Friedman test (Dunn test for multiple comparisons). Fractional occupancy and mean dwell times of each state at baseline and 60 minutes were compared with a paired t-test or Wilcoxon test.

## Results

### Barrel and Retrosplenial Areas Emerge as Key Regions of Functional Connectivity Alterations Following CSD

First, we looked at how functional connectivity among various cortical regions is impacted shortly after CSD. Since trigeminal nociceptive signals are relayed to contralateral cortical areas, our expectations were towards functional connectivity alterations on the hemisphere contralateral to CSD induction (i.e., the left hemisphere in our model). Indeed, several cortical regions on that side, including the barrel field, which receives nociceptive signals through the trigeminal sensory system ^11^, and the retrosplenial area, part of the default mode network ^71–73^, showed distinct alterations in functional connectivity and network topology, 30 and 60 minutes after CSD. Specifically, the contralateral barrel field showed increased connectivity with contralateral somatosensory forelimb and secondary motor regions, along with the ipsilateral barrel field (Figure 4). On the other hand, the contralateral barrel field showed increased anticorrelation with the ipsilateral retrosplenial area, while the contralateral retrosplenial region had increased anticorrelation with bilateral secondary motor regions, and decreased anticorrelation with the contralateral hindlimb area 60 minutes after CSD (Figure 4). We also noticed several alterations in connectivity of ipsilateral and contralateral secondary motor areas (Figure 4, 5, Supplementary Figures 2, 3, 5). All these changes in functional connectivity after CSD in the control group were prevented by naproxen administration (Figures 4, 5).

Chronic amitriptyline-administered mice had decreased susceptibility to CSD with increased CSD thresholds (Figure 3C) as expected ^25^. We realized that the resting-state functional network responded differently to CSD in the amitriptyline-treated group. We did not detect an increase in connectivity of barrel cortices with other somatosensory areas, unlike the control group, while the contralateral retrosplenial area had increased anticorrelations with both secondary motor areas (also seen in controls) and both barrel fields (unlike controls) 60 minutes after CSD (Figure 4, 5). We also detected connectivity changes in the bilateral hindlimb region and secondary motor areas and the ipsilateral forelimb somatosensory cortex (Figures 4, 5, Supplementary Figure 3, 4).

**Figure 3.**
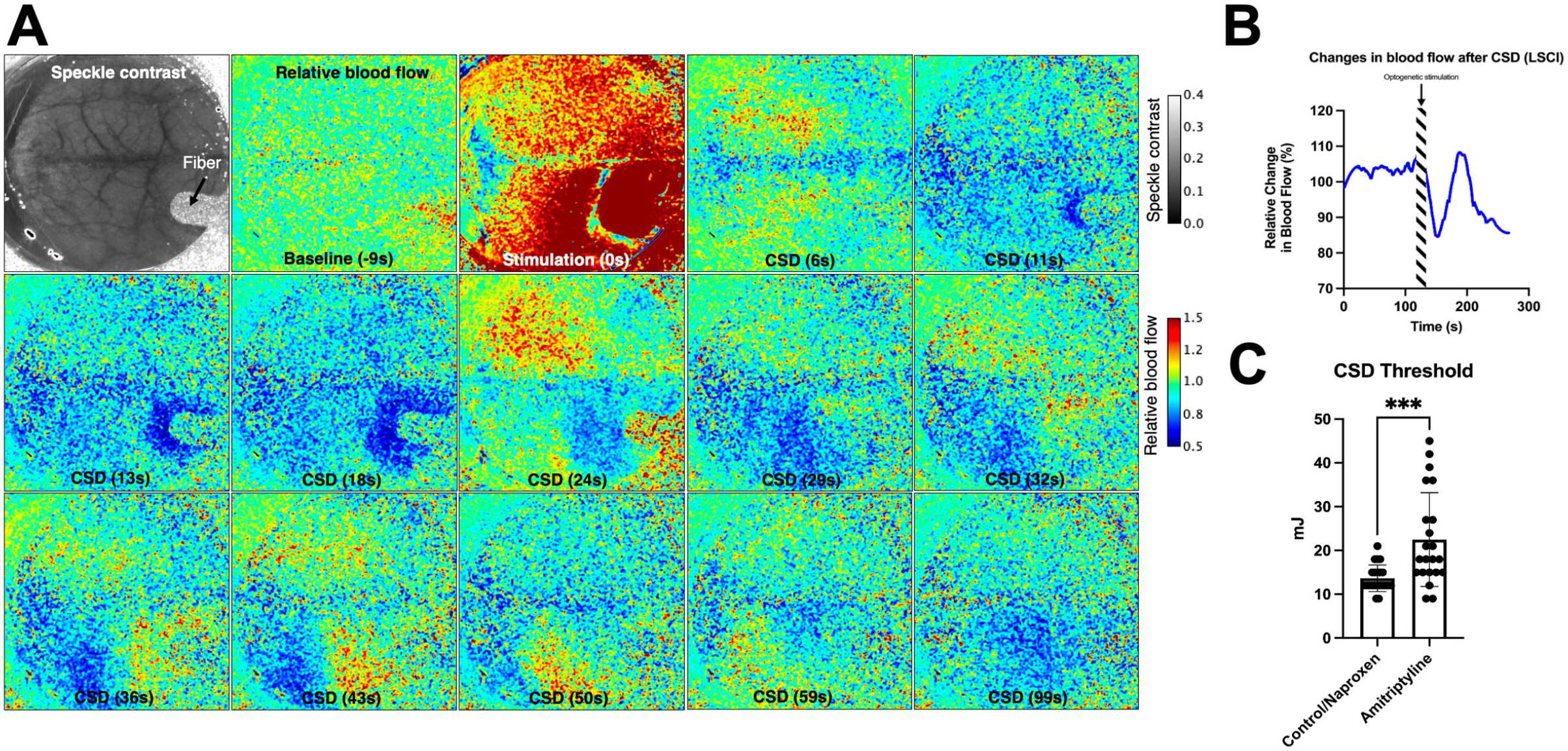
**(A)** Laser speckle contrast and relative blood flow maps as the CSD wave propagates unilaterally over the hemisphere. **(B)** Representative trace of relative blood flow changes in the ipsilateral cortex after CSD. **(C)** CSD thresholds of control/naproxen and chronic amitriptyline-administered mice. Panel (A) is reproduced from Solgun et al. ^26^ with permission.

**Figure 4.**
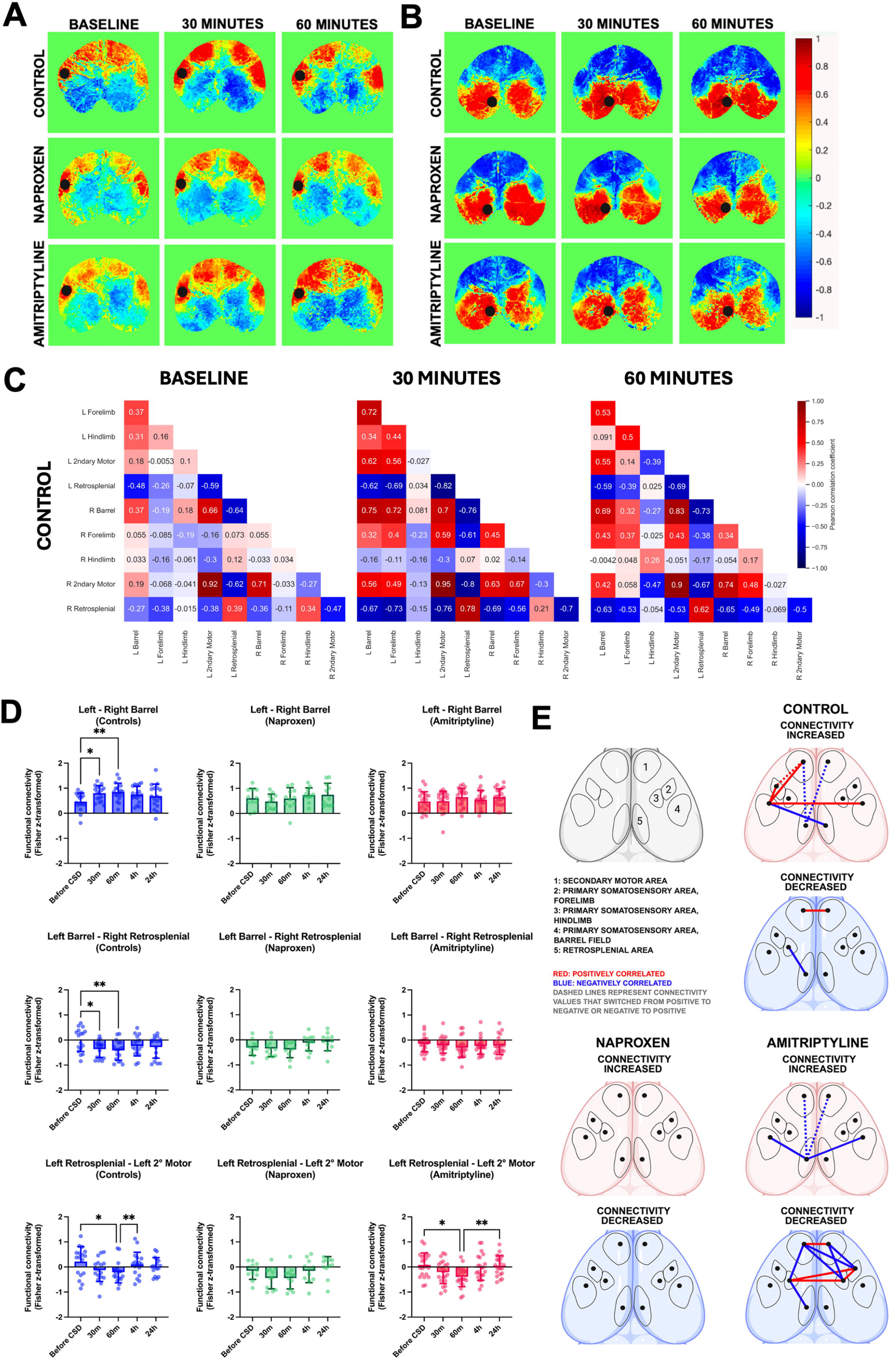
**(A-B)** Seed-based connectivity maps from representative animals with seeds placed at contralateral barrel **(A)** and contralateral retrosplenial **(B)** cortices of each group at baseline, 30 and 60 minutes after CSD. In each map, the correlation coefficients of the chosen seed to all other pixels are shown. **(C)** Correlation maps from a representative animal in the control group at baseline, 30 and 60 minutes after CSD. Correlation maps of representative animals from other groups can be found in Supplementary Figure 1B **(D)** A subset of functional connectivity alterations of each group are shown, remaining graphs can be found in Supplementary Figures 2-4. **(E)** Functional connectivity alterations at 30 and/or 60 minutes are shown in 3 groups, upper panel reflects connectivity values increased at 30 and/or 60 minutes relative to baseline, while lower panel shows decreased values decreased at 30 and/or 60 minutes relative to baseline. Red lines show positively correlated, blue lines show negatively correlated values, dashed lines show values reversed from positive to negative or negative to positive compared to baseline. One-way ANOVA for repeated measures (Tukey test for multiple comparisons) or Friedman test (Dunn test for multiple comparisons). *P<0.05, **P<0.01, ***P<0.001.

**Figure 5.**
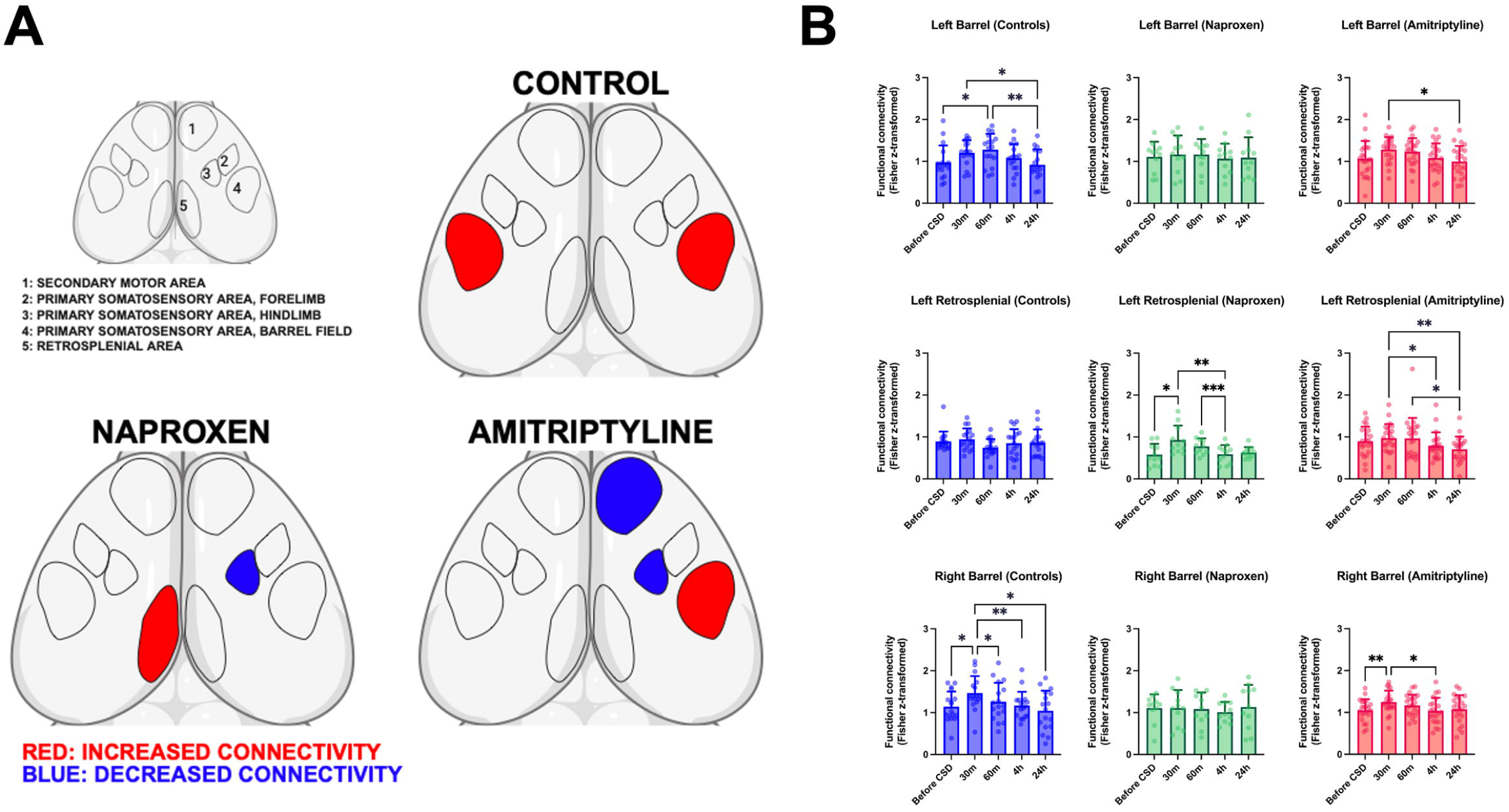
**(A)** Within-region functional connectivity alterations at 30 and/or 60 minutes are shown in all 3 groups. Red regions show increased, blue regions show decreased correlations at 30 and/or 60 minutes, relative to baseline values. **(B)** A subset of functional connectivity changes of each group, at different time points before and after CSD induction are shown. Remaining comparisons can be found in Supplementary Figure 5. Vertical bars indicate mean ± standard deviation. One-way ANOVA for repeated measures (Tukey test for multiple comparisons) or Friedman test (Dunn test for multiple comparisons). *P<0.05, **P<0.01, ***P<0.001.

Next, we looked at within-region (internal) connectivity, a measure of the functional connectivity of different seeds within the same region of interest with each other, as an indicator of coordinated neuronal activity in a particular cortical area. At 30 and 60 minutes after CSD, control mice displayed increased connectivity in the barrel field bihemispherically, a finding which was absent in the naproxen group (Figure 5A, 5B), but still marginally present in the amitriptyline group (Figure 5B). On the other hand, the naproxen group showed increased connectivity in the contralateral retrosplenial cortex at 30 minutes (Figure 5). Internal connectivity of contralateral retrosplenial cortex was also slightly higher at 30-60 minutes in the amitriptyline group, but not significantly different from the baseline (while still higher than 4-hour and 24-hour timepoints after CSD) (Figure 5B). These results indicate a similar within-region connectivity change in barrel cortices in both control and amitriptyline groups, while left retrosplenial cortex within-region connectivity showed a similar profile between naproxen and amitriptyline-treated groups. Amitriptyline-treated mice additionally showed decreased connectivity in the ipsilateral hindlimb somatosensory area (another similarity with naproxen-treated mice), and in the ipsilateral secondary motor cortex (Figure 5A, Supplementary Figure 5). These results indicate that both amitriptyline and naproxen change the connectivity of retrosplenial and somatosensory cortices, which could potentially indicate a headache-modulatory effect.

### Susceptibility to CSD does not alter baseline functional connectivity

We also wanted to investigate whether changes in CSD susceptibility were reflected in baseline functional connectivity patterns. We first divided animals into three groups based on their CSD thresholds: high ( ≥18 mJ), medium (15 mJ) and low (≤12 mJ) CSD threshold. Next, we compared the baseline connectivity values of high ( ≥18 mJ, n=17) and low (≤12 mJ, n=16) CSD threshold groups. We did not find any differences in within or between region connectivity values between groups after FDR correction (Supplementary Figure 6A). Next, we compared baseline connectivity values of amitriptyline (-) (control and naproxen groups) and amitriptyline (+) groups, which revealed no statistically significant differences (Supplementary Figure 6B).

### Barrel field contralateral to CSD induction plays a more prominent hub-like role after CSD

Next, we explored how the cortical functional architecture changes with CSD using a graph-theoretical approach to better assess the prior connectivity change findings at a network scale. In controls, eigenvector centrality and degree centrality of the barrel field contralateral to CSD increased at 60 minutes following CSD. PageRank centrality of the barrel field contralateral to CSD also displayed a near-significant increase at 60 minutes (One-way ANOVA for repeated measures, FDR-corrected p= 0.0567, Tukey test for multiple comparisons, p= 0.0170). Ipsilateral barrel field also displayed increased eigenvector centrality and degree centrality at 60 minutes compared to baseline (Figure 6A, 6D). Naproxen administration prevented the increase in all three centrality measures in barrel cortices (Figure 6B, 6E).

**Figure 6.**
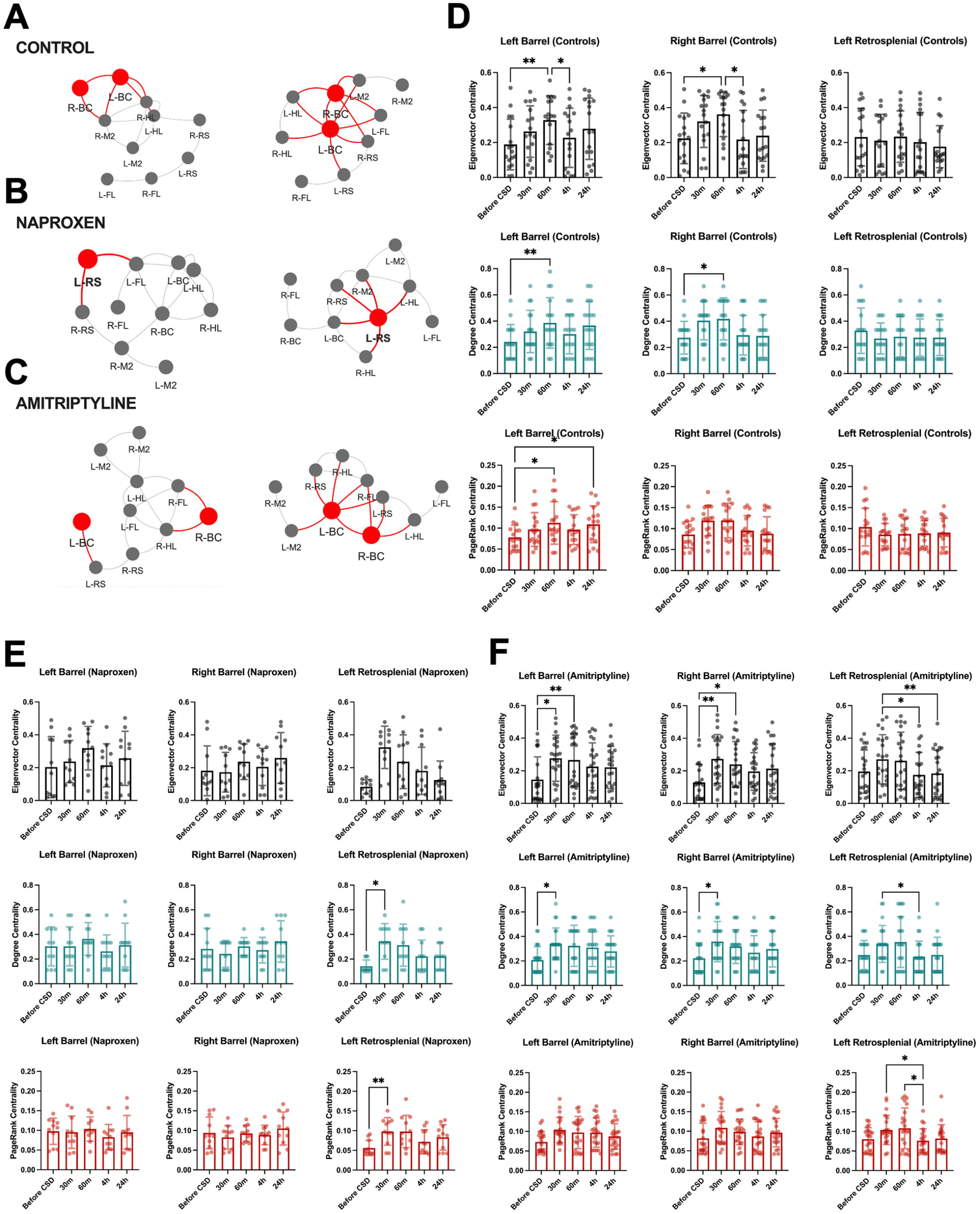
Graph-theoretical representation of brain network topology from representative mice from three groups mouse at baseline and after CSD **(A-C).** Centrality measures of barrel and contralateral retrosplenial areas contralateral to CSD in controls, naproxen, and amitriptyline groups **(D-F)**. L-BC: left (contralateral) barrel field, R-BC: right (ipsilateral) barrel field, L-HL: left hindlimb area, R-HL: right hindlimb area, L-FL: left forelimb area, R-FL: right forelimb area, L-M2: left secondary motor area, R-M2: right secondary motor area, L-RS: left retrosplenial area, R-RS: right retrosplenial area. Note that near-significant results include: R-BC degree centrality in the amitriptyline group (Friedman test, FDR-corrected p = 0.0586, Dunn test for multiple comparisons (Before CSD-30m), p = 0.0267), L-BC Pagerank centrality in the control group (One-way ANOVA for repeated measures, FDR-corrected p = 0.0567, Tukey test for multiple comparisons, (Before CSD-60m p = 0.0170, Before CSD-24h p = 0.0306), L-RS Pagerank centrality in the naproxen group (Friedman test, FDR-corrected p = 0.059, Dunn test for multiple comparisons, p = 0.0075), L-RS Pagerank centrality in the amitriptyline group (Friedman test, FDR-corrected p = 0.0575, Dunn test for multiple comparisons, (30m-4h (p = 0.0312), 60m-4h (p= 0.0312)). Vertical bars indicate mean ± standard deviation. One-way ANOVA for repeated measures (Tukey test for multiple comparisons) or Friedman test (Dunn test for multiple comparisons). *P<0.05, **P<0.01.

In the amitriptyline group, we detected a similar pattern in both contralateral and ipsilateral barrel fields, with degree centrality and eigenvector centrality of the contralateral barrel field increasing as early as 30 minutes (Figure 6C, 6F). Ipsilateral barrel field displayed an increase in eigenvector centrality and a near-significant increase in degree centrality (Friedman test, FDR-corrected p= 0.0586, Dunn test for multiple comparisons, p= 0.0267) after CSD.

In the naproxen group, the contralateral retrosplenial area showed increased degree centrality, with a near-significant increase in PageRank centrality (Friedman test, FDR-corrected p= 0.059, Dunn test for multiple comparisons, p=0.0075) following CSD at 30 minutes compared to baseline (Figure 6B, 6E). Centrality measures of the retrosplenial area did not increase in the control group. For the chronic amitriptyline group, although changes in centrality measures of left retrosplenial cortex did not reach statistical significance at 30-60 min after CSD, we noticed a tendency for increased centrality at these time points.

Additionally, hindlimb area ipsilateral to CSD, a region with multiple alterations in connectivity with other regions as well as decreased within region connectivity in the naproxen and chronic amitriptyline-treated groups, had decreased degree centrality in the naproxen group.

These findings, when evaluated together with the increased connectivity of the bilateral barrel field in the control group and contralateral retrosplenial field in the naproxen-administered group and possibly in the amitriptyline-treated group, once again brings up a potential role of these two regions in pain modulation in the resting-state functional network after CSD.

### Susceptibility to CSD does not alter baseline centrality measures

Next, we investigated whether changes in CSD susceptibility were reflected in baseline centrality measures. We again divided animals into three groups based on their CSD thresholds and compared the baseline connectivity values of high ( ≥18 mJ, n=17) and low (≤12 mJ, n=16) CSD threshold groups. We did not find any differences in any of the centrality measures between groups after FDR correction (Supplementary Figure 8A). We also compared baseline centrality measures values of amitriptyline (-) (control and naproxen groups) and amitriptyline (+) groups, which did not demonstrate any statistically significant differences (Supplementary Figure 8B).

### Mouse Grimace Scale scores indicate transient headache after CSD and are correlated with functional connectivity alterations in the barrel cortex in controls

Building on these findings, to evaluate and confirm the presence of headache and pain after CSD in our experimental approach, we used a modified version of the Mouse Grimace Scale (MGS), an established approach to evaluate pain from facial expressions in mice ^51,74–77^. The control group displayed increased modified MGS scores after CSD, indicating headache, while modified MGS scores in the naproxen-administered group did not differ from baseline (Figure 7A, 7B). Interestingly, amitriptyline administration did not prevent modified MGS increase after CSD, and modified MGS scores increased after CSD in the amitriptyline group as well, similar to controls (Figure 7A, 7B).

**Figure 7.**
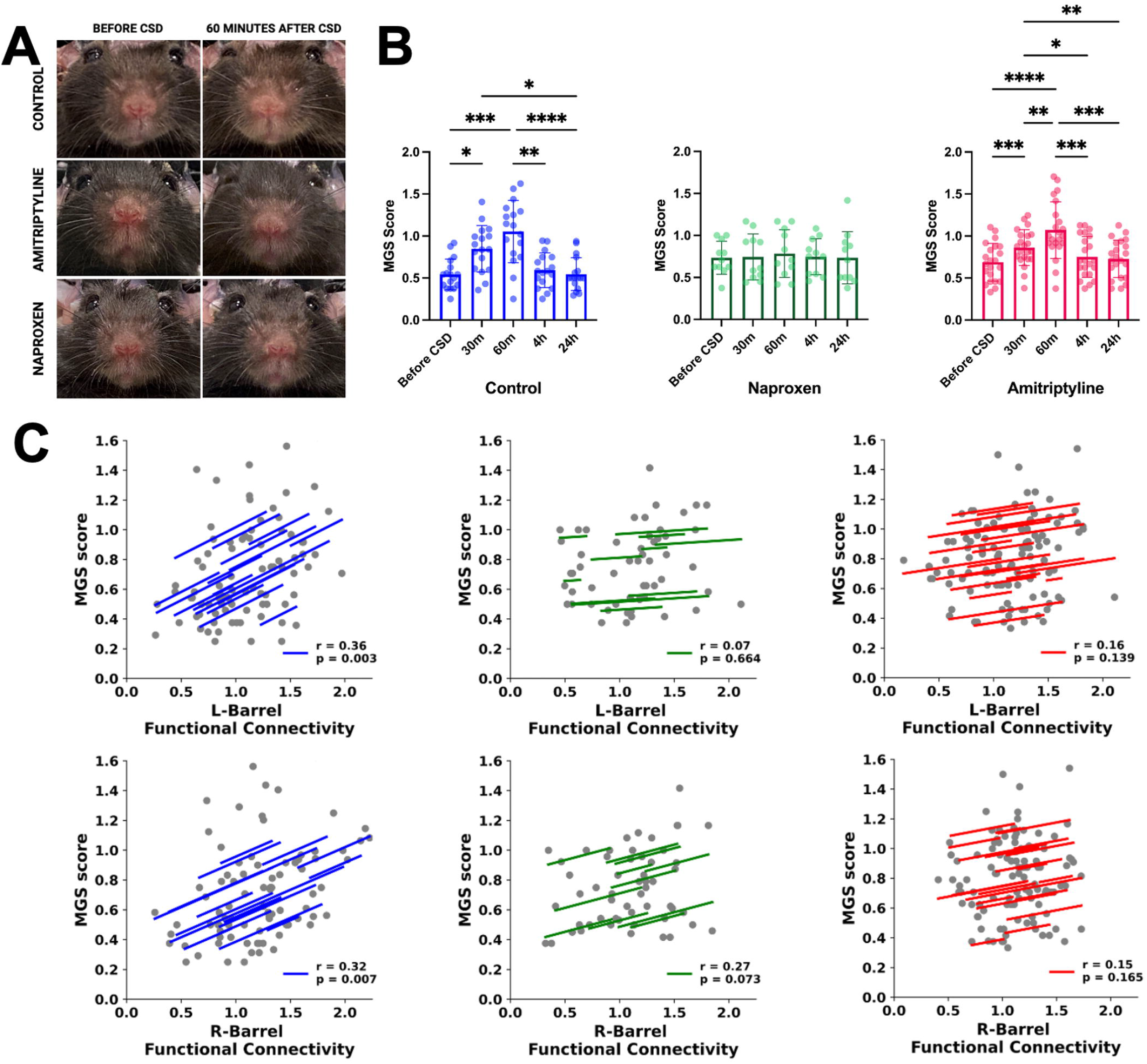
**(A)** Photographs of representative mice from control, amitriptyline, and naproxen groups before and 60 minutes after CSD. **(B)** Modified MGS scores at baseline and after CSD in controls, naproxen-administered animals, and the amitriptyline group. **(C)** Repeated-measures correlation plots of barrel cortices for each experimental group. Vertical bars indicate mean ± standard deviation. One-way ANOVA for repeated measures (Tukey test for multiple comparisons) or Friedman test (Dunn test for multiple comparisons). *P<0.05, **P<0.01, ***P<0.001, ****P<0.0001.

We next investigated whether the functional connectivity alterations described above were correlated with pain severity. We used repeated measures correlation, which investigates whether there is a reliable association between functional connectivity values and modified MGS scores for each animal using all five imaging sessions from baseline to 24 hours after CSD. In control animals, repeated measures correlation analysis showed significant correlation between modified MGS scores and within-region connectivity of the left and right barrel cortex (Figure 7C). There was no significant correlation between barrel cortex connectivity and modified MGS scores in naproxen or amitriptyline groups (Figure 7C).

### Dynamic functional connectivity analysis reveals potential CSD and pain-modulated states

Recent studies have highlighted that functional connectivity is not fixed across time, and regions have fluctuating connectivity patterns that cannot be revealed with a static connectivity approach, leading to a growing recognition of dynamic functional connectivity in the field ^68,78–84^. Dynamic or time-varying functional connectivity uses sliding-window correlations across an imaging session to reveal transient functional connectivity changes that cannot be captured by static functional connectivity approaches. We computed dynamic functional connectivity values between all 10 regions of interest, in addition to internal dynamic functional connectivity values within all 10 regions of interest, using sliding-window correlations.

State-specific measures from k-means clustering of dynamic functional connectivity, such as fractional occupancy and mean dwell time, demonstrated the effect of CSD on the network across all groups, in addition to the impact of pain in groups that were not treated by naproxen. Principal component analysis of clustered data can be visualized in Figure 8A. Pearson correlation coefficients of state centroids can be seen in Figure 8B. A decrease in fractional occupancy and mean dwell time of State 3 in all animals suggested that State 3 is possibly a state modulated by CSD, regardless of headache. Meanwhile, while fractional occupancy and mean dwell time of State 1 increased in control and amitriptyline groups, we observed no statistically significant change in the naproxen group, suggesting that State 1 is potentially a pain-modulated state (Figure 8C, 8D, 8E). State 1 is a state with high positive functional connectivity between bihemispheric barrel cortices and bihemispheric secondary motor cortices. While State 3, a potentially CSD-modulated state, has positive correlations between the left retrosplenial cortex and bihemispheric secondary motor cortices, State 1 displays anticorrelations. In addition, in State 1, in contrast with State 3, the left barrel cortex is anti-correlated with the right retrosplenial cortex, and the left retrosplenial cortex is weakly correlated with the left hindlimb area, further demonstrating that these two states have distinct functional connectivity features.

**Figure 8.**
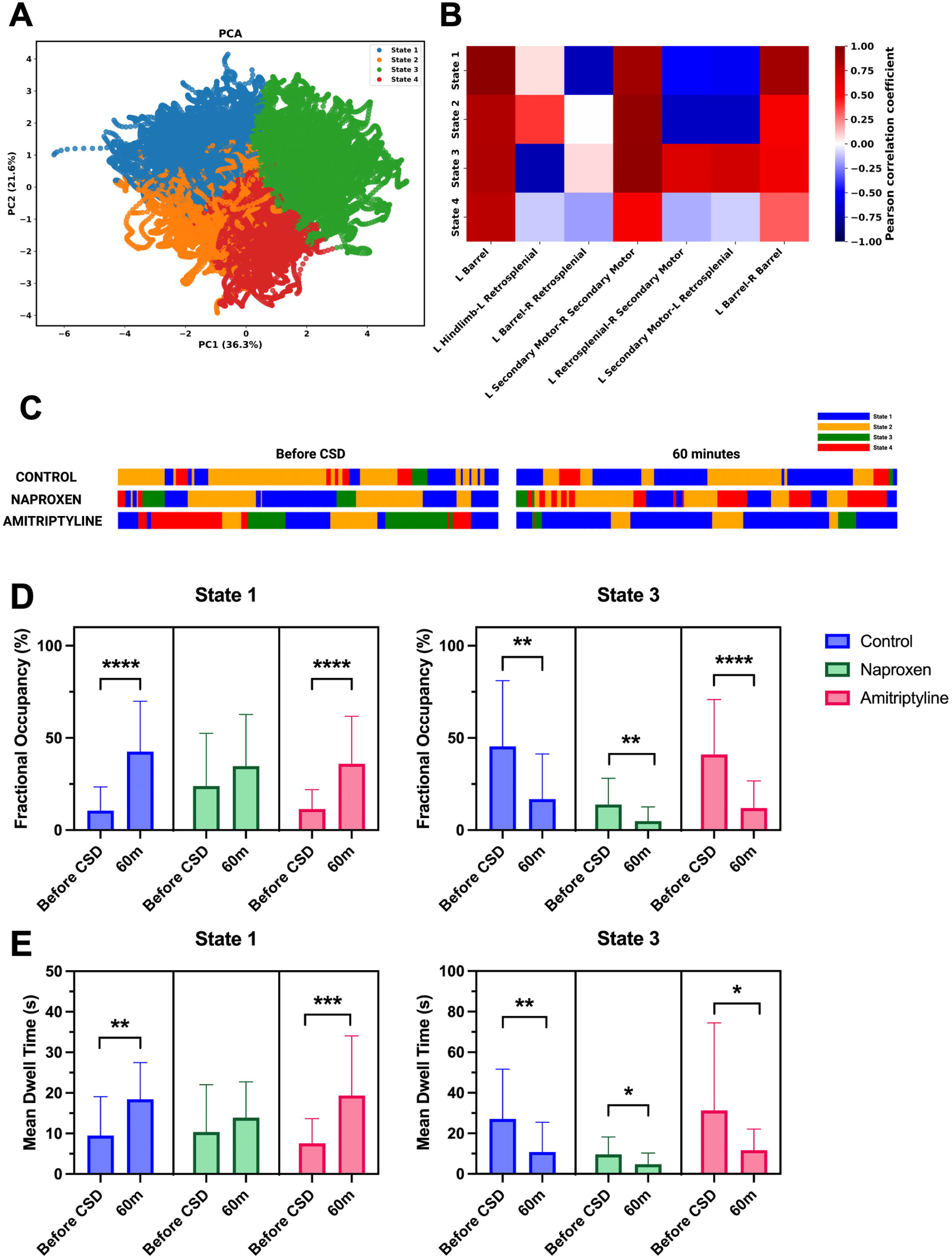
**(A)** Principal component analysis of dynamic functional connectivity data clustered into states with k-means clustering. **(B)** Pearson correlation coefficients of state centroids. **(C)** Representative **e**vent plots of states throughout an imaging session before and 60 minutes after CSD from representative animals from each of the groups. **(D)** Fractional occupancy of States 1 and 3 before and 60 minutes after CSD in all three groups. **(E)** Mean dwell time of States 1 and 3 before and 60 minutes after CSD in all three groups. Vertical bars indicate mean ± standard deviation. Paired t-test or Wilcoxon test. *P<0.05, **P<0.01, ***P<0.001, ****P<0.0001.

### Classification performance of a Support Vector Machine classifier trained on dynamic functional connectivity values

In view of our findings, which point to distinct alterations in the functional organization of the resting-state cortical network following CSD, possibly due to cortical pain modulation, we next utilized a support vector machine (SVM) to better assess findings from a classification perspective. Dynamic functional connectivity results in better classifier accuracy than static functional connectivity ^83,85^; therefore, to train our model, we chose dynamic functional connectivity values at baseline and 60 minutes. Since control mice display the unaltered cortical pain modulation response to CSD, which possibly reflects a functional pain signature, we used altered functional connectivity values (combined between and within region connectivity values) from the control group as features, resulting in 7 features in total (Figure 9A) ^79,81,86–88^. Four features were selected during cross-validation based on ANOVA F-values (shown in green in Figure 9A). ROI pairs involving contralateral barrel, retrosplenial, and secondary motor areas were the most discriminative features and chosen by the model during cross-validation. SVM trained on control data had 73.1% accuracy and 79.3% AUC in differentiating functional connectivity values at baseline and 60 minutes, based only on resting-state dynamic functional connectivity data. While solely using dynamic functional connectivity may only lead to moderate classifier performance, we chose this approach since our priority was to evaluate potential imaging correlates of trigeminal pain processing using functional connectivity values. When timepoint labels (baseline, 60 minutes) were randomly shuffled, classification accuracy decreased to 48.6%. AUC, accuracy, and F1 scores with confusion matrices can be seen in

**Figure 9.**
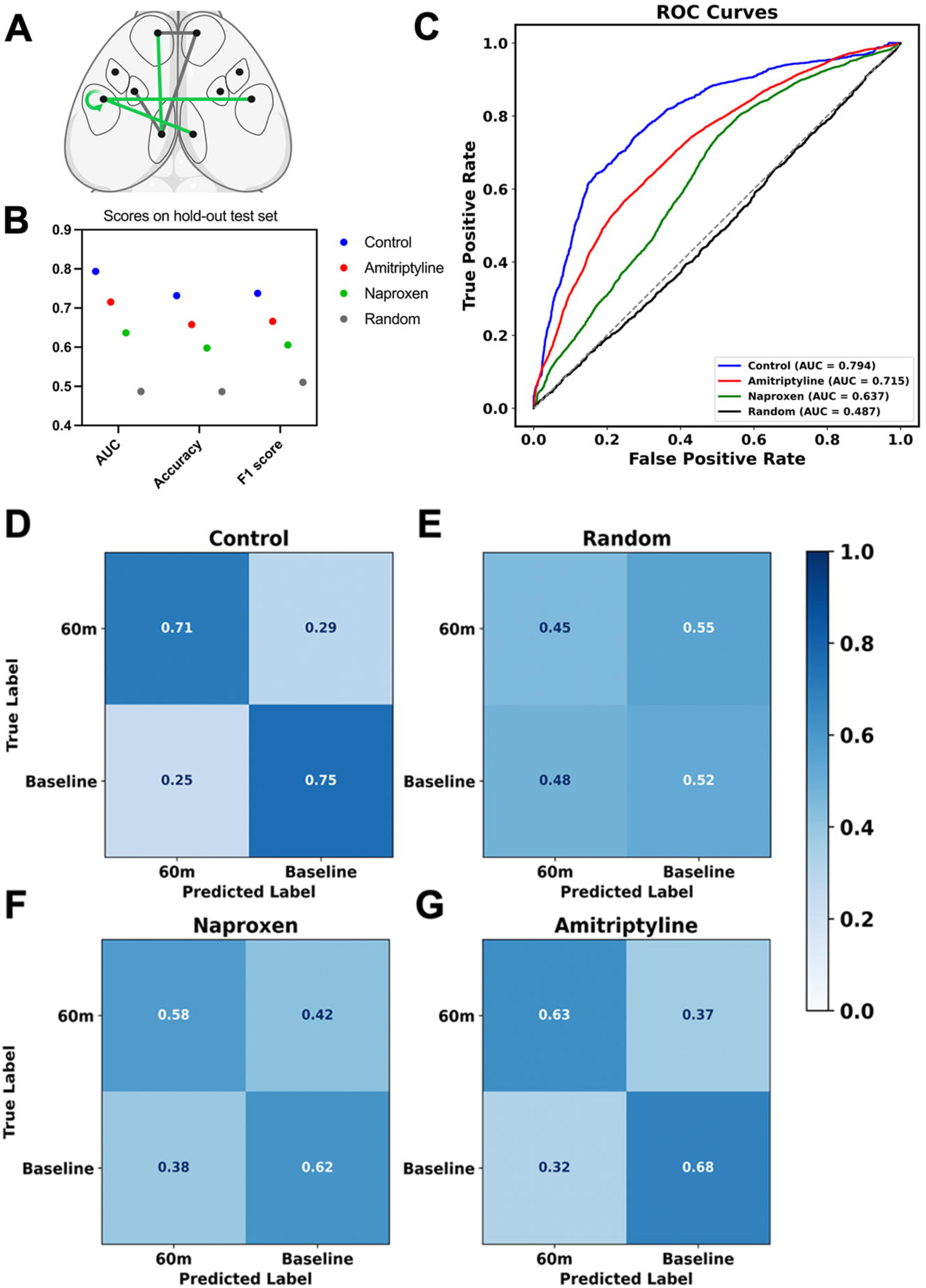
**(A)** Connectivity values that show significant change at 60 minutes in controls relative to baseline; these were used as features for cross-validation. Features selected inside cross-validation are shown in green. **(B)** Area under the curve (AUC), accuracy, and F1 scores for control, naproxen, and amitriptyline groups on the test set. **(C)** Receiver operating characteristic (ROC) curves for control, naproxen, and amitriptyline groups on the test set. **(D-G)** Confusion matrices for controls **(D)** and randomly shuffled labels in controls **(E)**, naproxen **(F),** and amitriptyline **(G)** groups.

Figure 9 (See Methods for model details). Additionally, cross-prediction with the model trained on the control group performed poorly when tested on the naproxen-administered or chronic amitriptyline-administered group (accuracy scores at 59.8% and 65.7%, respectively), indicating distinct alterations in how the resting-state cortical network of these groups responds to CSD and trigeminal pain modulation on the cortical surface. Notably, classification performance exhibited a graded pattern across groups, with the highest performance in the control group, the lowest performance in the naproxen group, and an intermediate performance in the amitriptyline group. This gradient suggests that the classifier is sensitive to the degree of similarity to control-driven dynamic functional connectivity patterns, a potential “functional pain signature”. The intermediate accuracy in the amitriptyline group indicates partial overlap in dynamic functional connectivity patterns, consistent with the static functional connectivity results, whereas the reduced performance in the naproxen group is consistent with the lack of statistically significant alterations in static functional connectivity between regions of interest in this group. Overall, the graded decline in cross-prediction performance suggests that the classifier captures control-derived dynamic functional connectivity structure that is only partially preserved in a similar group such as amitriptyline and largely absent in the naproxen-administered group.

## Discussion

There are several experimental readouts of trigeminovascular activation, the underlying mechanism of migraine headache, such as middle meningeal artery dilation, increased c-fos expression in the trigeminocervical complex, electrophysiological activation of the trigeminal ganglion and the trigeminal nucleus caudalis, and blood neuropeptide level alterations, all of which have been shown to be affected after CSD ^32,89–94^, the putative cause of migraine aura. While these experimental techniques are useful for experimental migraine research, it has to be kept in mind that these are indirect indicators of the process, while trigeminal afferents ultimately project to the cerebral cortex as headache is perceived by the subject. It could be an influential tool if we could find evidence of headache-related cortical processing in the rodent brain. The primary aim of this work is to disclose the cortical representation of CSD-triggered headache in awake mice, following a minimally invasive approach. Our findings highlight the interplay between potential role players in the “pain matrix” ^95,96^, in the context of trigeminovascular activation following CSD in awake mice, and provide information about potential modulatory areas, via the effect of an acute abortive medication (like naproxen) or a chronic prophylactic medication (amitriptyline).

We demonstrate that CSD is associated with transient changes in resting-state functional connectivity and network-level reorganization on the cortical surface in a rodent model of migraine, under awake conditions. These alterations became maximal in about 60 minutes and started to normalize after 4 hours, mainly involving the barrel field in the control group, with increased centrality of the barrel field in the resting-state cortical network. Importantly, these changes are prevented by naproxen administration, suggesting that these changes are not due to nonspecific cortical activity modulation by CSD but rather related to inflammatory activation and possibly headache induction. The positive correlation of connectivity measures with modified MGS scores (i.e., headache severity) in the control group is also suggestive of this association. We additionally see a differential involvement of the contralateral retrosplenial cortex in the naproxen-administered group, with increased internal connectivity and centrality after CSD. On the other hand, amitriptyline, a medication used in migraine prophylaxis, increased the CSD threshold, altered how the resting-state cortical network responded to CSD, but did not change the baseline connectivity. Amitriptyline did not completely prevent the within-region connectivity and centrality of barrel fields after CSD and did not prevent the increased dominance of dynamic State-1, and this is in concordance with our finding that amitriptyline also did not prevent the increase in modified MGS after CSD. These data might provide clues about the mechanism of preventive action of amitriptyline for migraine; it mainly increases the CSD threshold, preventing aura, and does not fully prevent headache and connectivity changes in somatosensory areas once CSD is triggered. On the other hand, the increased involvement of contralateral retrosplenial cortex in the amitriptyline group, similar to the effect of acute naproxen administration, might still have a modulatory effect in post-CSD headache, perhaps via increasing the effectiveness of acute abortive medications. We did not test the effect of naproxen in the amitriptyline-treated group, but this is a potential area for future exploration.

As migraine is a disorder of sensory processing ^97^ increased connectivity of contralateral primary somatosensory cortex, specifically the barrel field, is consistent with its activation in response to nociceptive stimuli from downstream areas ^11,48–50,98,99^ and several studies showcase functional connectivity alterations of the sensorimotor network in migraine ^12,13,46,47,100^. Altered functional connectivity in subregions of the sensorimotor network and with other brain regions has been reported in migraine patients without aura in the interictal period, supporting previous reports indicating a dysfunction of multisensory integration in migraine ^101^. In an awake rat fMRI study using the inflammatory-soup model to induce headache, authors reported increased connectivity of the default mode network (DMN) with sensorimotor areas, increased connectivity of the somatosensory cortex with the sensorimotor network, and altered connectivity of the salience network with somatosensory areas ^102^.

On the other hand, functional alterations are not limited to the somatosensory cortex following CSD. Alterations in the retrosplenial cortex with analgesic administration are especially notable in our study, given that the retrosplenial cortex, one of the main areas in the DMN in mice ^71–73^, is a major region involved in pain perception, with a potential modulatory role in pain processing ^37–45,103^. Increased anti-correlation between the ipsilateral retrosplenial cortex and contralateral somatosensory cortex was detected in the control group after CSD, while this finding was not seen in the naproxen and amitriptyline groups. The retrosplenial cortex most likely functions as a “pain-integrator” of sensory, cognitive, and emotional aspects of pain in the network, and constitutes a part of the functional pain signature ^38,47,100,103,104^ along with a potential role in pain-related contextual memory ^41^. The role of retrosplenial cortex in analgesia/antinociception, possibly through descending pain inhibitory mechanisms ^39,40,42–44,105,106^, is in concordance with the observed alterations in this study, namely the increased centrality and within-region connectivity of the retrosplenial cortex in the naproxen-administered group. This is clinically supported as decreased functional connectivity of the cerebellum with DMN has been shown in episodic migraine patients without aura interictally, which was negatively correlated with migraine frequency ^107^.

Our classifier utilized dynamic resting-state functional connectivity values and demonstrated moderate classification performance to distinguish between baseline and post-CSD conditions. Classification performance was highest in controls, while cross-prediction accuracy in amitriptyline followed second, and cross-prediction accuracy in the naproxen group was lowest, signifying that the model was not able to distinguish between functional connectivity metrics before and after CSD in the naproxen group, consistent with the fact that the naproxen group revealed no difference in static functional connectivity between regions. It is worth noting that higher classification accuracy scores could be possibly achieved through combining multiple measures of functional connectivity, using graph-based measures, combined structural connectivity values, static and/or dynamic functional connectivity values, state-based measures after clustering, or clinical, behavioral, or demographic information ^108–110^. We refrained from combining behavioral readouts like MGS scores or network topology measures like centrality in our study since our aim was to uncover which of the functional connectivity features were key players in distinguishing the impact of CSD, in search of a functional pain “signature” of CSD and headache.

This study has several strengths. First is the minimally invasive skull optical clearing and CSD induction technique, coupled with awake imaging. Since anesthesia disrupts neurovascular coupling and functional connectivity patterns and alters CSD duration ^111–116^, our approach brings insight into how the natural state of the brain is impacted by cortical spreading depression and the resulting headache, and the mechanisms by which common abortive and prophylactic migraine medications might exert their effects. Additionally, compared to cross-sectional interictal clinical studies providing snapshots of network properties compared to healthy controls, this study also highlighted that network alterations were transient and, after a peak at 30 to 60 minutes, a return to baseline was present at 4 to 24 hours after CSD. To our knowledge, this is the only awake minimally invasive functional imaging study in rodents that investigates the effect of pain on the resting-state cortical network after CSD. Finally, we also implement dynamic functional connectivity measures, which could reveal the time-varying nature of pain perception and processing ^117^. Our results revealed potential CSD and pain-modulated states, providing insight into the dynamic nature of the network after CSD beyond a static functional connectivity approach. Our work may be valuable from a translational aspect to investigate future clinical targets in migraine. For instance, functional near-infrared spectroscopy (fNIRS) is an emerging field, and measurements from the superficial cortical surface can be utilized in further studies, for which studies examining the cortical surface, like ours, can act as a guide^118^.

Some limitations, on the other hand, should be considered. Our results are restricted to the cortical surface due to the nature of our experimental setup, and deeper cortical and subcortical structures that contribute to pain processing could not be investigated. In addition, our sample only had male mice since female mice may have inconsistent CSD thresholds that depend on the phase of the estrous cycle and may also have different thresholds than male mice ^119–121^. In addition, only undirected measures of connectivity, such as correlation, were studied. Because of the correlative nature of our findings, it is not known if these topological alterations are adaptive, compensatory, or simply an impact of pain on the network. Future studies utilizing directed measures, such as effective connectivity between these regions, will be critical for further understanding of top-down and bottom-up pain modulation after CSD.

## Conclusion

In the wake of CSD, there is early mesoscale functional reorganization of cortical surface-level networks previously implicated in pain perception and processing. These alterations are evident shortly after CSD and return to baseline at or before 24 hours, possibly reflecting trigeminal pain modulation on the cortical surface after CSD. Together, our findings show that CSD is associated with reorganization of mesoscale cortical dynamics, characterized by network-level alterations and altered state metrics. Mesoscale static and dynamic functional connectivity approaches in rodents provide a sensitive framework to understand cortical pain modulation with CSD and migraine.

## Data Availability

Raw and processed datasets, as well as the codes and the trained Support Vector Machine used during the current study, are available from the corresponding author upon reasonable request.

## Acknowledgements

We thank Kaan Kutay Özmen, MD, PhD for design and 3D-printing of the headplates with screwable caps, that were crucial for mouse fixation, exposure of both hemispheres, keeping the clearing material in place, protecting the window from dust and preventing optogenetic activation outside experimental sessions. These plates were also used in our prior work^26^. We thank Sreekanth Kura, PhD, from Boston University Neurophotonics Center, for providing the functional connectivity analysis code and his assistance. Figures were created with BioRender.

## Author Contributions

B.S. designed and performed the experiments, acquired, processed, and analyzed the data, and drafted the manuscript. B.S. and A.B.V. performed MGS analyses. B.D.D. and H.K. contributed to experiments and animal-related procedures. E.D. supervised data analysis and SVM classifier applications. Ş.E.E. conceived the study, designed and supervised the experiments, analyzed and interpreted the data, drafted and critically revised the manuscript.

## References

1. Olesen J, Bes A, Kunkel R, et al. The International Classification of Headache Disorders, 3rd edition (beta version). Cephalalgia. 2013;33(9):629–808. doi:10.1177/0333102413485658

2. Russel MB, Olesen J. A nosographic analysis of the migraine aura in a general population. Brain. 1996;119 ( Pt 2)(2):355–361. doi:10.1093/BRAIN/119.2.355

3. Hadjikhani N, Sanchez Del Rio M, Wu O, et al. Mechanisms of migraine aura revealed by functional MRI in human visual cortex. Proc Natl Acad Sci U S A. 2001;98(8):4687. doi:10.1073/PNAS.071582498

4. Lipton RB, Bigal ME, Diamond M, Freitag F, Reed ML, Stewart WF. Migraine prevalence, disease burden, and the need for preventive therapy. Neurology. 2007;68(5):343–349. doi:10.1212/01.wnl.0000252808.97649.21

5. Bloudek LM, Stokes M, Buse DC, et al. Cost of healthcare for patients with migraine in five European countries: results from the International Burden of Migraine Study (IBMS). J Headache Pain. 2012;13(5):361–378. doi:10.1007/s10194-012-0460-7

6. Blumenfeld A, Varon S, Wilcox T, et al. Disability, HRQoL and resource use among chronic and episodic migraineurs: Results from the International Burden of Migraine Study (IBMS). Cephalalgia. 2011;31(3):301–315. doi:10.1177/0333102410381145

7. Steiner TJ, Stovner LJ, Vos T, Jensen R, Katsarava Z. Migraine is first cause of disability in under 50s: will health politicians now take notice? J Headache Pain. 2018;19(1):17. doi:10.1186/s10194-018-0846-2

8. Ashina M, Terwindt GM, Al-Karagholi MAM, et al. Migraine: disease characterisation, biomarkers, and precision medicine. The Lancet. 2021;397(10283):1496–1504. doi:10.1016/S0140-6736(20)32162-0

9. Skorobogatykh K, van Hoogstraten WS, Degan D, et al. Functional connectivity studies in migraine: what have we learned? J Headache Pain. 2019;20(1):108. doi:10.1186/s10194-019-1047-3

10. Lo Buono V, Bonanno L, Corallo F, et al. Functional connectivity and cognitive impairment in migraine with and without aura. J Headache Pain. 2017;18(1). doi:10.1186/S10194-017-0782-6

11. Cui Y, Toyoda H, Sako T, et al. A voxel-based analysis of brain activity in high-order trigeminal pathway in the rat induced by cortical spreading depression. Neuroimage. 2015;108:17–22. doi:10.1016/j.neuroimage.2014.12.047

12. Amin FM, Hougaard A, Magon S, et al. Change in brain network connectivity during PACAP38-induced migraine attacks. Neurology. 2016;86(2):180–187. doi:10.1212/WNL.0000000000002261

13. Zhang J, Su J, Wang M, et al. The sensorimotor network dysfunction in migraineurs without aura: a resting-state fMRI study. J Neurol. 2017;264(4):654–663. doi:10.1007/s00415-017-8404-4

14. Coppola G, Di Renzo A, Tinelli E, et al. Resting state connectivity between default mode network and insula encodes acute migraine headache. Cephalalgia. 2018;38(5):846–854. doi:10.1177/0333102417715230

15. Chong CD, Gaw N, Fu Y, Li J, Wu T, Schwedt TJ. Migraine classification using magnetic resonance imaging resting-state functional connectivity data. Cephalalgia. 2017;37(9):828–844. doi:10.1177/0333102416652091

16. Hu S, Hao Z, Li M, et al. Resting-state abnormalities in functional connectivity of the default mode network in migraine: A meta-analysis. Front Neurosci. 2023;17. doi:10.3389/fnins.2023.1136790

17. Harriott AM, Takizawa T, Chung DY, Chen SP. Spreading depression as a preclinical model of migraine. J Headache Pain. 2019;20(1):45. doi:10.1186/s10194-019-1001-4

18. Ayata C, Shin HK, Salomone S, et al. Pronounced Hypoperfusion during Spreading Depression in Mouse Cortex. Journal of Cerebral Blood Flow & Metabolism. 2004;24(10):1172–1182. doi:10.1097/01.WCB.0000137057.92786.F3

19. Charles AC, Baca SM. Cortical spreading depression and migraine. Nat Rev Neurol. 2013;9(11):637-644. doi:10.1038/nrneurol.2013.192

20. Leao AAP. SPREADING DEPRESSION OF ACTIVITY IN THE CEREBRAL CORTEX. 101152/jn194476359. 1944;7(6):359–390. doi:10.1152/JN.1944.7.6.359

21. Vinogradova L V, Suleymanova EM, Medvedeva TM. Transient loss of interhemispheric functional connectivity following unilateral cortical spreading depression in awake rats. Cephalalgia. 2021;41(3):353–365. doi:10.1177/0333102420970172

22. Lachinova DA, Smirnova MP, Pavlova I V., Sysoev I V., Vinogradova L V. Transient destabilization of interhemispheric functional connectivity induced by spreading depolarization. Network Neuroscience. Published online July 18, 2024:1–37. doi:10.1162/netn_a_00405

23. Li B, Zhou F, Luo Q, Li P. Altered resting-state functional connectivity after cortical spreading depression in mice. Neuroimage. 2012;63(3):1171–1177. doi:10.1016/j.neuroimage.2012.08.024

24. White BR, Bauer AQ, Snyder AZ, Schlaggar BL, Lee JM, Culver JP. Imaging of Functional Connectivity in the Mouse Brain. PLoS One. 2011;6(1):e16322. doi:10.1371/journal.pone.0016322

25. Ayata C, Jin H, Kudo C, Dalkara T, Moskowitz MA. Suppression of cortical spreading depression in migraine prophylaxis. Ann Neurol. 2006;59(4):652–661. doi:10.1002/ana.20778

26. Solgun B, Dönmez-Demir B, Bahadır-Varol A, Karataş H, Erdener ŞE. Choosing a skull clearing technique for chronic mesoscopic optical imaging in awake mice. Neurophotonics. 2026;13(02). doi:10.1117/1.NPh.13.2.025006

27. Kyweriga M, Sun J, Wang S, Kline R, Mohajerani MH. A Large Lateral Craniotomy Procedure for Mesoscale Wide-field Optical Imaging of Brain Activity. J Vis Exp. 2017;(123). doi:10.3791/52642

28. Zhao YJ, Yu TT, Zhang C, et al. Skull optical clearing window for in vivo imaging of the mouse cortex at synaptic resolution. Light Sci Appl. 2017;7(2):17153–17153. doi:10.1038/lsa.2017.153

29. Li D, Hu Z, Zhang H, et al. A Through-Intact-Skull (TIS) chronic window technique for cortical structure and function observation in mice. eLight. 2022;2(1):15. doi:10.1186/s43593-022-00022-2

30. Zhang C, Liu C, Feng W. A Long-Term Clearing Cranial Window for Longitudinal Imaging of Cortical and Calvarial Ischemic Injury through the Intact Skull. Advanced Science. 2022;9(17). doi:10.1002/advs.202105893

31. Houben T, Loonen IC, Baca SM, et al. Optogenetic induction of cortical spreading depression in anesthetized and freely behaving mice. Journal of Cerebral Blood Flow & Metabolism. 2017;37(5):1641–1655. doi:10.1177/0271678X16645113

32. Karatas H, Erdener SE, Gursoy-Ozdemir Y, et al. Spreading Depression Triggers Headache by Activating Neuronal Panx1 Channels. Science (1979). 2013;339(6123):1092–1095. doi:10.1126/science.1231897

33. Kura S, Xie H, Fu B, Ayata C, Boas DA, Sakadžić S. Intrinsic optical signal imaging of the blood volume changes is sufficient for mapping the resting state functional connectivity in the rodent cortex. J Neural Eng. 2018;15(3):035003. doi:10.1088/1741-2552/aaafe4

34. Allen Reference Atlas – Mouse Brain. Available from atlas.brain-map.org.

35. Wang Q, Ding SL, Li Y, et al. The Allen Mouse Brain Common Coordinate Framework: A 3D Reference Atlas. Cell. 2020;181(4):936–953.e20. doi:10.1016/j.cell.2020.04.007

36. Rob Campbell (2025). AllenAtlasTopDown (https://github.com/Zapit-Optostim/AllenAtlasTopDown/releases/tag/v0.1.4), GitHub. Retrieved October 29, 2025.

37. Qiu Y, Lian YN, Wu C, Liu L, Zhang C, Li XY. Coordination between midcingulate cortex and retrosplenial cortex in pain regulation. Front Mol Neurosci. 2024;17. doi:10.3389/fnmol.2024.1405532

38. Barrière DA, Hamieh AM, Magalhães R, et al. Structural and functional alterations in the retrosplenial cortex following neuropathic pain. Pain. 2019;160(10):2241–2254. doi:10.1097/j.pain.0000000000001610

39. Reis GM, Dias QM, Silveira JWS, Del Vecchio F, Garcia-Cairasco N, Prado WA. Antinociceptive Effect of Stimulating the Occipital or Retrosplenial Cortex in Rats. J Pain. 2010;11(10):1015–1026. doi:10.1016/j.jpain.2010.01.269

40. Rossaneis AC, Reis GM, Prado WA. Stimulation of the occipital or retrosplenial cortex reduces incision pain in rats. Pharmacol Biochem Behav. 2011;100(2):220–227. doi:10.1016/j.pbb.2011.08.024

41. Hao S, Xue M, Chen QY, et al. Supraspinal facilitation of painful stimuli by glutamatergic innervation from the retrosplenial to the anterior cingulate cortex. PLoS Biol. 2025;23(1):e3003011. doi:10.1371/journal.pbio.3003011

42. Reis GM, Rossaneis AC, Silveira JWS, Dias QM, Prado WA. Stimulation-Produced Analgesia From the Occipital or Retrosplenial Cortex of Rats Involves Serotonergic and Opioid Mechanisms in the Anterior Pretectal Nucleus. J Pain. 2011;12(5):523–530. doi:10.1016/j.jpain.2010.10.006

43. Reis GM, Rossaneis AC, Silveira JWS, Prado WA. μ1- and 5-HT1-dependent mechanisms in the anterior pretectal nucleus mediate the antinociceptive effects of retrosplenial cortex stimulation in rats. Life Sci. 2012;90(23-24):950–955. doi:10.1016/j.lfs.2012.04.023

44. Reis GM, Fais RS, Prado WA. The antinociceptive effect of stimulating the retrosplenial cortex in the rat tail-flick test but not in the formalin test involves the rostral anterior cingulate cortex. Pharmacol Biochem Behav. 2015;131:112–118. doi:10.1016/j.pbb.2015.02.006

45. Vann SD, Aggleton JP, Maguire EA. What does the retrosplenial cortex do? Nat Rev Neurosci. 2009;10(11):792–802. doi:10.1038/nrn2733

46. Zhang Y, Liu N, Wang Z, et al. Impaired Inter-Hemispheric Functional Connectivity during Resting State in Female Patients with Migraine. Brain Sci. 2022;12(11):1505. doi:10.3390/brainsci12111505

47. Da Silva JT, Seminowicz DA. Neuroimaging of pain in animal models: a review of recent literature. Pain Rep. 2019;4(4):e732. doi:10.1097/PR9.0000000000000732

48. Ishikawa T, Murata K, Okuda H, et al. Pain-related neuronal ensembles in the primary somatosensory cortex contribute to hyperalgesia and anxiety. iScience. 2023;26(4):106332. doi:10.1016/j.isci.2023.106332

49. Xiao Z, Martinez E, Kulkarni PM, et al. Cortical Pain Processing in the Rat Anterior Cingulate Cortex and Primary Somatosensory Cortex. Front Cell Neurosci. 2019;13. doi:10.3389/fncel.2019.00165

50. Jin QQ, Wu GQ, Peng WW, Xia XL, Hu L, Iannetti GD. Somatotopic Representation of Second Pain in the Primary Somatosensory Cortex of Humans and Rodents. The Journal of Neuroscience. 2018;38(24):5538–5550. doi:10.1523/JNEUROSCI.3654-17.2018

51. Langford DJ, Bailey AL, Chanda ML, et al. Coding of facial expressions of pain in the laboratory mouse. Nat Methods. 2010;7(6):447–449. doi:10.1038/nmeth.1455

52. Duffy SS, Perera CJ, Makker PGS, Lees JG, Carrive P, Moalem-Taylor G. Peripheral and Central Neuroinflammatory Changes and Pain Behaviors in an Animal Model of Multiple Sclerosis. Front Immunol. 2016;7. doi:10.3389/fimmu.2016.00369

53. Whittaker AL, Liu Y, Barker TH. Methods Used and Application of the Mouse Grimace Scale in Biomedical Research 10 Years on: A Scoping Review. Animals. 2021;11(3):673. doi:10.3390/ani11030673

54. Bakdash JZ, Marusich LR. Repeated Measures Correlation. Front Psychol. 2017;8. doi:10.3389/fpsyg.2017.00456

55. Vallat R. Pingouin: statistics in Python. J Open Source Softw. 2018;3(31):1026. doi:10.21105/joss.01026

56. Kazeminejad A, Sotero RC. The Importance of Anti-correlations in Graph Theory Based Classification of Autism Spectrum Disorder. Front Neurosci. 2020;14. doi:10.3389/fnins.2020.00676

57. Zhan L, Jenkins LM, Wolfson OE, et al. The significance of negative correlations in brain connectivity. Journal of Comparative Neurology. 2017;525(15):3251–3265. doi:10.1002/cne.24274

58. Chen G, Chen G, Xie C, Li SJ. Negative Functional Connectivity and Its Dependence on the Shortest Path Length of Positive Network in the Resting-State Human Brain. Brain Connect. 2011;1(3):195–206. doi:10.1089/brain.2011.0025

59. Centeno EGZ, Moreni G, Vriend C, Douw L, Santos FAN. A hands-on tutorial on network and topological neuroscience. Brain Struct Funct. 2022;227(3):741–762. doi:10.1007/s00429-021-02435-0

60. Scharwächter L, Schmitt FJ, Pallast N, Fink GR, Aswendt M. Network analysis of neuroimaging in mice. Neuroimage. 2022;253:119110. doi:10.1016/j.neuroimage.2022.119110

61. Alexander-Bloch AF, Gogtay N, Meunier D, et al. Disrupted Modularity and Local Connectivity of Brain Functional Networks in Childhood-Onset Schizophrenia. Front Syst Neurosci. 2010;4. doi:10.3389/fnsys.2010.00147

62. Hagmann P, Cammoun L, Gigandet X, et al. Mapping the Structural Core of Human Cerebral Cortex. PLoS Biol. 2008;6(7):e159. doi:10.1371/journal.pbio.0060159

63. Fornito A, Zalesky A, Bullmore ET. Fundamentals of Brain Network Analysis. Elsevier/Academic Press; 2016.

64. Rubinov M, Sporns O. Complex network measures of brain connectivity: Uses and interpretations. Neuroimage. 2010;52(3):1059–1069. doi:10.1016/j.neuroimage.2009.10.003

65. Hagberg AA, Schult DA, Swart PJ. Exploring network structure, dynamics, and function using NetworkX. In: Varoquaux G, Vaught T, Millman J, eds. Proceedings of the 7th Python in Science Conference (SciPy2008). 2008:11–15.

66. Leonardi N, Van De Ville D. On spurious and real fluctuations of dynamic functional connectivity during rest. Neuroimage. 2015;104:430–436. doi:10.1016/j.neuroimage.2014.09.007

67. Pedregosa F, Varoquaux G, Gramfort A, et al. Scikit-learn: Machine Learning in Python. Journal of Machine Learning Research. 2011;12:2825–2830.

68. Allen EA, Damaraju E, Plis SM, Erhardt EB, Eichele T, Calhoun VD. Tracking Whole-Brain Connectivity Dynamics in the Resting State. Cerebral Cortex. 2014;24(3):663–676. doi:10.1093/cercor/bhs352

69. Genovese CR, Lazar NA, Nichols T. Thresholding of Statistical Maps in Functional Neuroimaging Using the False Discovery Rate. Neuroimage. 2002;15(4):870–878. doi:10.1006/nimg.2001.1037

70. Benjamini Y, Hochberg Y. Controlling the False Discovery Rate: A Practical and Powerful Approach to Multiple Testing. Journal of the Royal Statistical Society: Series B (Methodological). 1995;57(1):289–300. doi:10.1111/J.2517-6161.1995.TB02031.X

71. Lu H, Zou Q, Gu H, Raichle ME, Stein EA, Yang Y. Rat brains also have a default mode network. Proceedings of the National Academy of Sciences. 2012;109(10):3979–3984. doi:10.1073/pnas.1200506109

72. Upadhyay J, Baker SJ, Chandran P, et al. Default-Mode-Like Network Activation in Awake Rodents. PLoS One. 2011;6(11):e27839. doi:10.1371/journal.pone.0027839

73. Whitesell JD, Liska A, Coletta L, et al. Regional, Layer, and Cell-Type-Specific Connectivity of the Mouse Default Mode Network. Neuron. 2021;109(3):545–559.e8. doi:10.1016/j.neuron.2020.11.011

74. Dehghani A, Schenke M, van Heiningen SH, Karatas H, Tolner EA, van den Maagdenberg AMJM. Optogenetic cortical spreading depolarization induces headache-related behaviour and neuroinflammatory responses some prolonged in familial hemiplegic migraine type 1 mice. J Headache Pain. 2023;24(1):96. doi:10.1186/s10194-023-01628-8

75. Song X, Chen Z, Zhu H, et al. Transcutaneous Auricular Vagus Nerve Stimulation Alleviates Headache Symptoms in Migraine Model Mice by the Locus Coeruleus/Noradrenergic System: An Experimental Study in a Mouse Model of Migraine. Biomedicines. 2026;14(1):96. doi:10.3390/biomedicines14010096

76. Tzeng HR, Lee MT, Fan PC, et al. α6GABAA Receptor Positive Modulators Alleviate Migraine-like Grimaces in Mice via Compensating GABAergic Deficits in Trigeminal Ganglia. Neurotherapeutics. 2021;18(1):569–585. doi:10.1007/s13311-020-00951-1

77. Eller OC, Yang X, Fuentes IM, et al. Voluntary Wheel Running Partially Attenuates Early Life Stress-Induced Neuroimmune Measures in the Dura and Evoked Migraine-Like Behaviors in Female Mice. Front Physiol. 2021;12. doi:10.3389/fphys.2021.665732

78. Vergara VM, Mayer AR, Kiehl KA, Calhoun VD. Dynamic functional network connectivity discriminates mild traumatic brain injury through machine learning. Neuroimage Clin. 2018;19:30–37. doi:10.1016/j.nicl.2018.03.017

79. Lee MJ, Park BY, Cho S, Park H, Kim ST, Chung CS. Dynamic functional connectivity of the migraine brain: a resting-state functional magnetic resonance imaging study. Pain. 2019;160(12):2776–2786. doi:10.1097/j.pain.0000000000001676

80. Chen W, Hu H, Wu Q, et al. Altered Static and Dynamic Interhemispheric Resting-State Functional Connectivity in Patients With Thyroid-Associated Ophthalmopathy. Front Neurosci. 2021;15. doi:10.3389/fnins.2021.799916

81. Niu H, Li W, Wang G, et al. Performances of whole-brain dynamic and static functional connectivity fingerprinting in machine learning-based classification of major depressive disorder. Front Psychiatry. 2022;13. doi:10.3389/fpsyt.2022.973921

82. Rashid B, Arbabshirani MR, Damaraju E, et al. Classification of schizophrenia and bipolar patients using static and dynamic resting-state fMRI brain connectivity. Neuroimage. 2016;134:645–657. doi:10.1016/j.neuroimage.2016.04.051

83. Jin C, Jia H, Lanka P, et al. Dynamic brain connectivity is a better predictor of PTSD than static connectivity. Hum Brain Mapp. 2017;38(9):4479–4496. doi:10.1002/hbm.23676

84. Dumkrieger G, Chong CD, Ross K, Berisha V, Schwedt TJ. Static and dynamic functional connectivity differences between migraine and persistent post-traumatic headache: A resting-state magnetic resonance imaging study. Cephalalgia. 2019;39(11):1366–1381. doi:10.1177/0333102419847728

85. Nie W, Zeng W, Yang J, Zhao L, Shi Y. Classification of Migraine Using Static Functional Connectivity Strength and Dynamic Functional Connectome Patterns: A Resting-State fMRI Study. Brain Sci. 2023;13(4):596. doi:10.3390/brainsci13040596

86. Li M, Zhang S, Chu F, et al. Abnormal Static and Dynamic Functional Connectivity in Tension-Type Headache: A Support Vector Machine Analysis. J Neurosci Res. 2025;103(6). doi:10.1002/jnr.70057

87. Ichesco E, Peltier SJ, Mawla I, et al. Prediction of Differential Pharmacologic Response in Chronic Pain Using Functional Neuroimaging Biomarkers and a Support Vector Machine Algorithm: An Exploratory Study. Arthritis & Rheumatology. 2021;73(11):2127–2137. doi:10.1002/art.41781

88. Zhu J, Jiao Y, Chen R, Wang XH, Han Y. Aberrant dynamic and static functional connectivity of the striatum across specific low-frequency bands in patients with autism spectrum disorder. Psychiatry Res Neuroimaging. 2023;336:111749. doi:10.1016/j.pscychresns.2023.111749

89. Erdener SE, Dalkara T. Modelling headache and migraine and its pharmacological manipulation. Br J Pharmacol. 2014;171(20):4575–4594. doi:10.1111/bph.12651

90. Moskowitz M, Nozaki K, Kraig R. Neocortical spreading depression provokes the expression of c-fos protein-like immunoreactivity within trigeminal nucleus caudalis via trigeminovascular mechanisms. The Journal of Neuroscience. 1993;13(3):1167–1177. doi:10.1523/JNEUROSCI.13-03-01167.1993

91. Zhang X, Levy D, Noseda R, Kainz V, Jakubowski M, Burstein R. Activation of Meningeal Nociceptors by Cortical Spreading Depression: Implications for Migraine with Aura. Journal of Neuroscience. 2010;30(26):8807–8814. doi:10.1523/JNEUROSCI.0511-10.2010

92. Zhang X, Levy D, Kainz V, Noseda R, Jakubowski M, Burstein R. Activation of central trigeminovascular neurons by cortical spreading depression. Ann Neurol. 2011;69(5):855–865. doi:10.1002/ana.22329

93. Kaag Rasmussen M, Møllgård K, Bork PAR, et al. Trigeminal ganglion neurons are directly activated by influx of CSF solutes in a migraine model. Science (1979). 2024;385(6704):80–86. doi:10.1126/science.adl0544

94. Schain AJ, Melo-Carrillo A, Stratton J, Strassman AM, Burstein R. CSD-Induced Arterial Dilatation and Plasma Protein Extravasation Are Unaffected by Fremanezumab: Implications for CGRP’s Role in Migraine with Aura. The Journal of Neuroscience. 2019;39(30):6001–6011. doi:10.1523/JNEUROSCI.0232-19.2019

95. Legrain V, Iannetti GD, Plaghki L, Mouraux A. The pain matrix reloaded. Prog Neurobiol. 2011;93(1):111–124. doi:10.1016/j.pneurobio.2010.10.005

96. Thompson SJ, Bushnell MC. Rodent functional and anatomical imaging of pain. Neurosci Lett. 2012;520(2):131–139. doi:10.1016/j.neulet.2012.03.015

97. Goadsby PJ, Holland PR, Martins-Oliveira M, Hoffmann J, Schankin C, Akerman S. Pathophysiology of Migraine: A Disorder of Sensory Processing. Physiol Rev. 2017;97(2):553–622. doi:10.1152/physrev.00034.2015

98. Hjornevik T, Jacobsen LM, Qu H, Bjaalie JG, Gjerstad J, Willoch F. Metabolic plasticity in the supraspinal pain modulating circuitry after noxious stimulus-induced spinal cord LTP. Pain. 2008;140(3):456–464. doi:10.1016/j.pain.2008.09.029

99. Yue L, Bao C, Zhang L, et al. Neuronal mechanisms of nociceptive-evoked gamma-band oscillations in rodents. Neuron. 2025;113(5):769–784.e6. doi:10.1016/j.neuron.2024.12.011

100. Jia Z, Chen X, Tang W, Zhao D, Yu S. Atypical functional connectivity between the anterior cingulate cortex and other brain regions in a rat model of recurrent headache. Mol Pain. 2019;15. doi:10.1177/1744806919842483

101. Qin Z, Su J, He XW, et al. Disrupted functional connectivity between sub-regions in the sensorimotor areas and cortex in migraine without aura. J Headache Pain. 2020;21(1):47. doi:10.1186/s10194-020-01118-1

102. Becerra L, Bishop J, Barmettler G, Kainz V, Burstein R, Borsook D. Brain network alterations in the inflammatory soup animal model of migraine. Brain Res. 2017;1660:36–46. doi:10.1016/j.brainres.2017.02.001

103. Wik G, Fischer H, Finer B, Bragee B, Kristianson M, Fredrikson M. RETROSPENIAL CORTICAL DEACTIVATION DURING PAINFUL STIMULATION OF FIBROMYALGIC PATIENTS. International Journal of Neuroscience. 2006;116(1):1–8. doi:10.1080/00207450690962208

104. Seminowicz DA, Jiang L, Ji Y, Xu S, Gullapalli RP, Masri R. Thalamocortical Asynchrony in Conditions of Spinal Cord Injury Pain in Rats. The Journal of Neuroscience. 2012;32(45):15843–15848. doi:10.1523/JNEUROSCI.2927-12.2012

105. Zhao W, Liu SL, Lin SS, Zhang Y, Yu C. Astrocytic P2X7 receptor in retrosplenial cortex drives electroacupuncture analgesia. Purinergic Signal. 2025;21(4):523–532. doi:10.1007/s11302-024-10043-w

106. Lin WY, Chu WH, Chao THH, Sun WZ, Yen CT. Longitudinal FDG-PET scan study of brain changes in mice with cancer-induced bone pain and after morphine analgesia. Mol Pain. 2019;15. doi:10.1177/1744806919841194

107. Ke J, Yu Y, Zhang X, et al. Functional Alterations in the Posterior Insula and Cerebellum in Migraine Without Aura: A Resting-State MRI Study. Front Behav Neurosci. 2020;14. doi:10.3389/fnbeh.2020.567588

108. Hsiao FJ, Chen WT, Liu HY, et al. Altered brainstem–cortex activation and interaction in migraine patients: somatosensory evoked EEG responses with machine learning. J Headache Pain. 2024;25(1):185. doi:10.1186/s10194-024-01892-2

109. Zhang Q, Wu Q, Zhang J, et al. Discriminative Analysis of Migraine without Aura: Using Functional and Structural MRI with a Multi-Feature Classification Approach. PLoS One. 2016;11(9):e0163875. doi:10.1371/journal.pone.0163875

110. Messina R, Sudre CH, Wei DY, Filippi M, Ourselin S, Goadsby PJ. Biomarkers of Migraine and Cluster Headache: Differences and Similarities. Ann Neurol. 2023;93(4):729–742. doi:10.1002/ana.26583

111. Boveroux P, Vanhaudenhuyse A, Bruno MA, et al. Breakdown of within- and between-network Resting State Functional Magnetic Resonance Imaging Connectivity during Propofol-induced Loss of Consciousness. Anesthesiology. 2010;113(5):1038–1053. doi:10.1097/ALN.0b013e3181f697f5

112. Xie H, Chung DY, Kura S, et al. Differential effects of anesthetics on resting state functional connectivity in the mouse. Journal of Cerebral Blood Flow & Metabolism. 2020;40(4):875–884. doi:10.1177/0271678X19847123

113. Aksenov DP, Li L, Miller MJ, Iordanescu G, Wyrwicz AM. Effects of Anesthesia on BOLD Signal and Neuronal Activity in the Somatosensory Cortex. Journal of Cerebral Blood Flow & Metabolism. 2015;35(11):1819–1826. doi:10.1038/jcbfm.2015.130

114. Bukhari Q, Schroeter A, Cole DM, Rudin M. Resting State fMRI in Mice Reveals Anesthesia Specific Signatures of Brain Functional Networks and Their Interactions. Front Neural Circuits. 2017;11. doi:10.3389/fncir.2017.00005

115. Grandjean J, Schroeter A, Batata I, Rudin M. Optimization of anesthesia protocol for resting-state fMRI in mice based on differential effects of anesthetics on functional connectivity patterns. Neuroimage. 2014;102:838–847. doi:10.1016/j.neuroimage.2014.08.043

116. Masamoto K, Fukuda M, Vazquez A, Kim S. Dose-dependent effect of isoflurane on neurovascular coupling in rat cerebral cortex. European Journal of Neuroscience. 2009;30(2):242–250. doi:10.1111/j.1460-9568.2009.06812.x

117. Kucyi A, Davis KD. The dynamic pain connectome. Trends Neurosci. 2015;38(2):86-95. doi:10.1016/j.tins.2014.11.006

118. Ferrari M, Quaresima V. A brief review on the history of human functional near-infrared spectroscopy (fNIRS) development and fields of application. Neuroimage. 2012;63(2):921–935. doi:10.1016/j.neuroimage.2012.03.049

119. Brennan KC, Romero-Reyes M, López Valdés HE, Arnold AP, Charles AC. Reduced threshold for cortical spreading depression in female mice. Ann Neurol. 2007;61(6):603–606. doi:10.1002/ana.21138

120. Kudo C, Harriott AM, Moskowitz MA, Waeber C, Ayata C. Estrogen modulation of cortical spreading depression. J Headache Pain. 2023;24(1):62. doi:10.1186/s10194-023-01598-x

121. Ebine T, Toriumi H, Shimizu T, et al. Alterations in the threshold of the potassium concentration to evoke cortical spreading depression during the natural estrous cycle in mice. Neurosci Res. 2016;112:57–62. doi:10.1016/j.neures.2016.06.001

