## Supplemental Figures for "Mesoscale Functional Reorganization of Cortical Networks After Cortical Spreading Depression"

**Supplementary Figure 1 (A)** Seed selection for between-regions (left) and within-region (right) connectivity analyses. **(B)** Correlation maps of representative animals from amitriptyline and naproxen groups at baseline, 30 and 60 minutes after CSD.

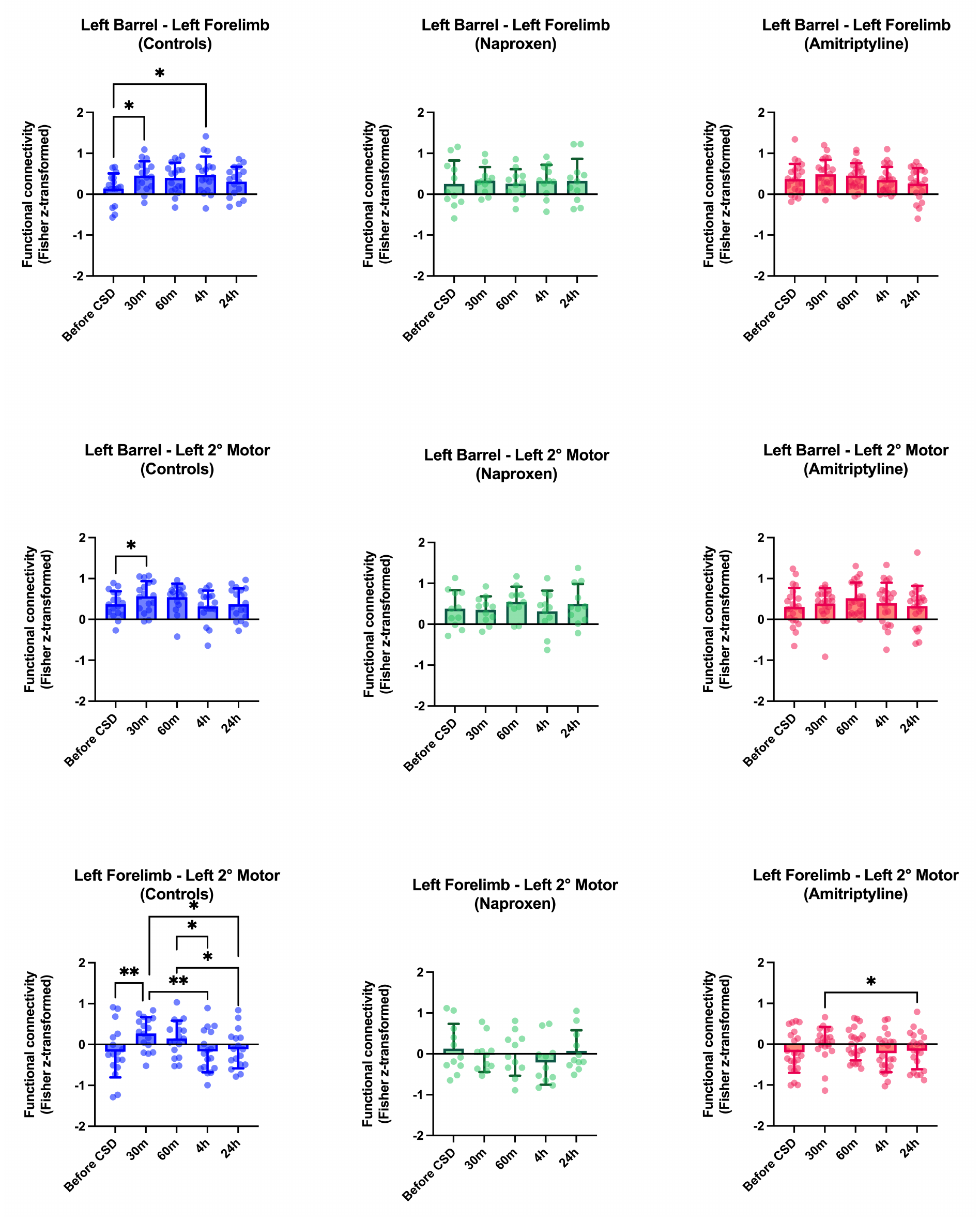

**Supplementary Figure 2** Functional connectivity alterations of each group, remaining alterations statistically significant in only the control group are presented. One-way ANOVA for repeated measures (Tukey test for multiple comparisons) or Friedman test (Dunn test for multiple comparisons). *P<0.05, **P<0.01.

**Supplementary Figure 3** Functional connectivity alterations of each group, remaining alterations statistically significant in control and amitriptyline groups are presented. One-way ANOVA for repeated measures (Tukey test for multiple comparisons) or Friedman test (Dunn test for multiple comparisons). *P<0.05, **P<0.01, ***P<0.001.

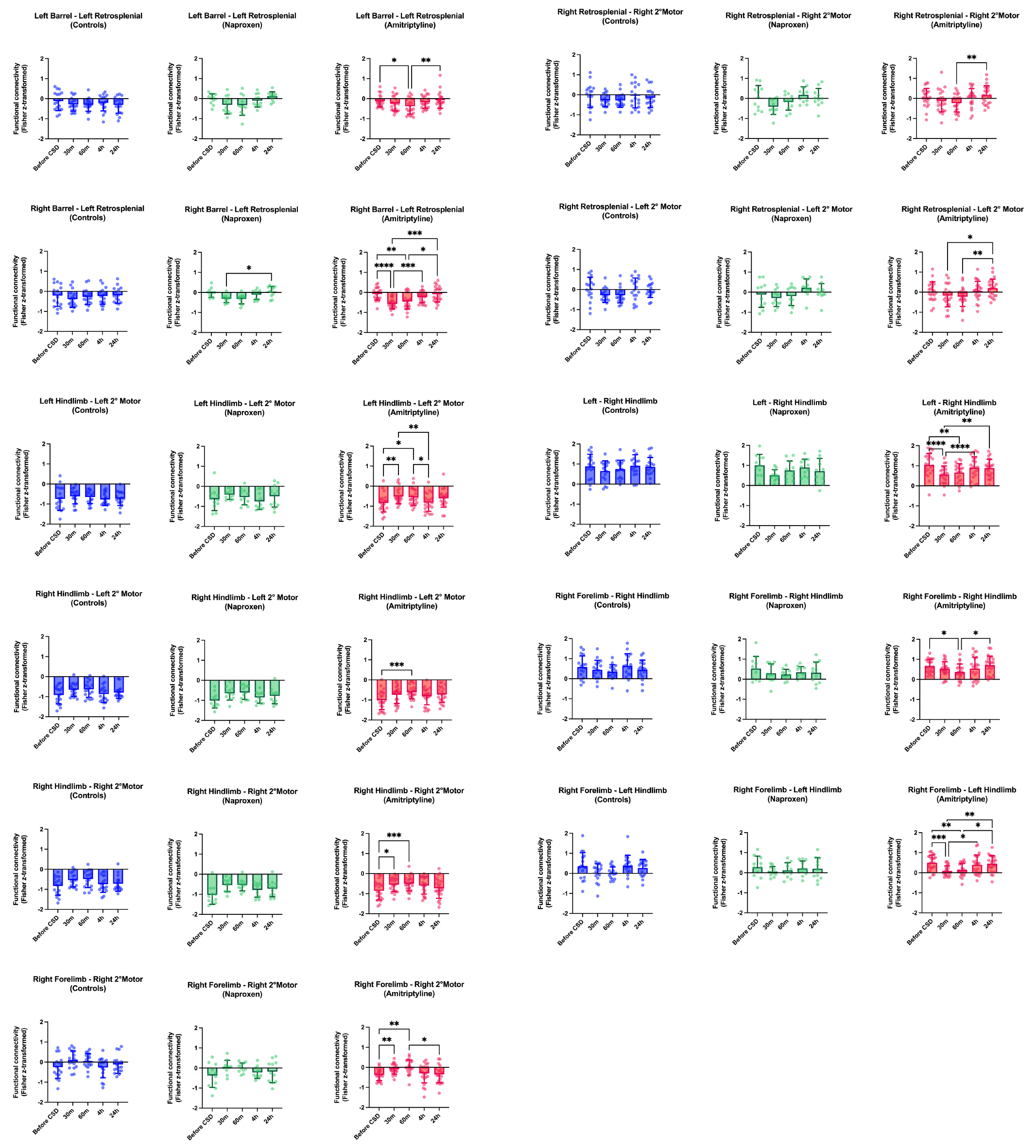

**Supplementary Figure 4** Functional connectivity alterations of each group, remaining alterations statistically significant in only the amitriptyline groups are presented. One-way ANOVA for repeated measures (Tukey test for multiple comparisons) or Friedman test (Dunn test for multiple comparisons). *P<0.05, **P<0.01, ***P<0.001, ****P<0.0001.

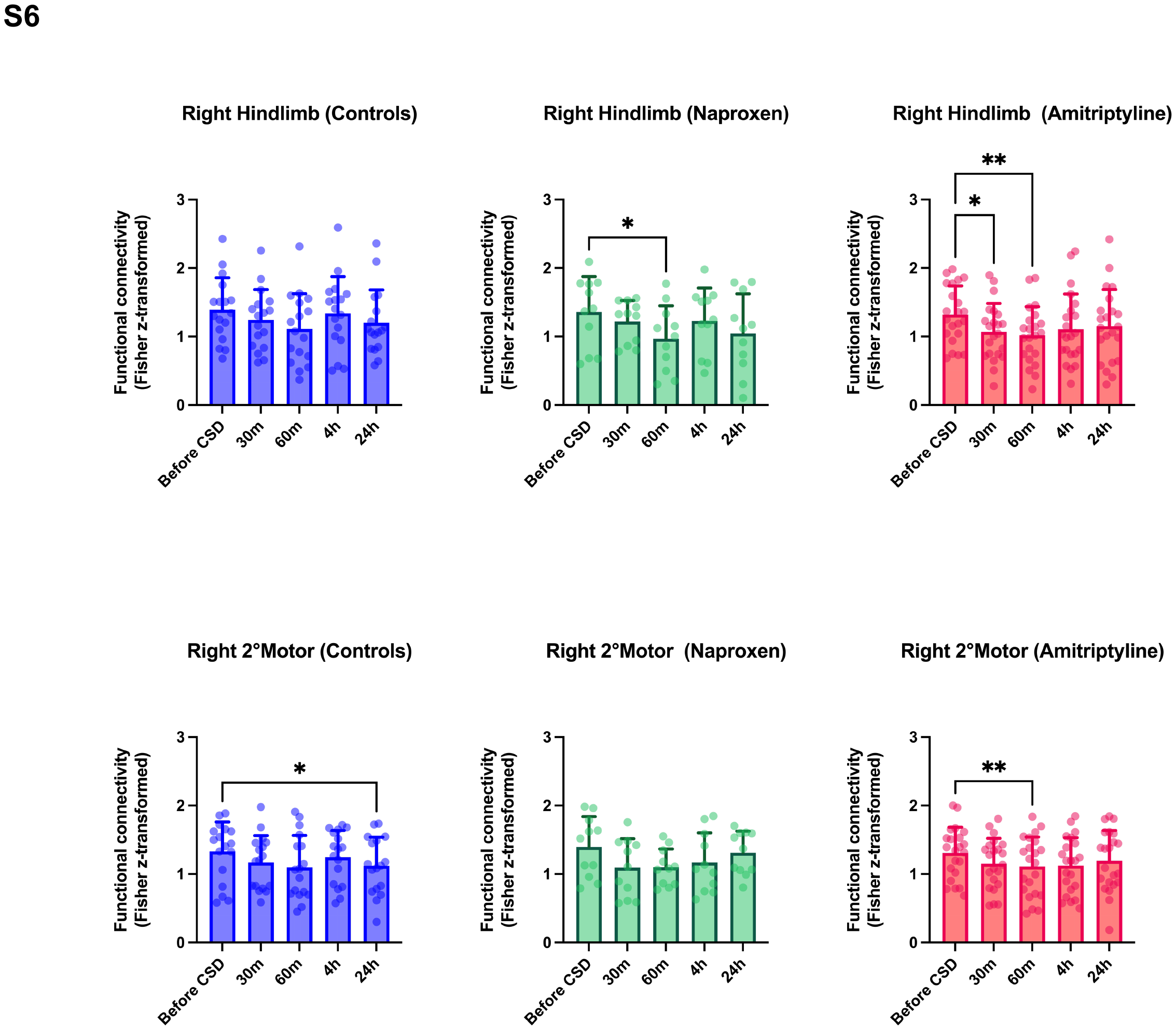

**Supplementary Figure 5** Remaining statistically-significant within-region functional connectivity alterations of each group. One-way ANOVA for repeated measures (Tukey test for multiple comparisons) or Friedman test (Dunn test for multiple comparisons). *P<0.05, **P<0.01.

**Supplementary Figure 6 (A)** Baseline functional connectivity comparisons of animals with high and low CSD threshold. **(B)** Baseline functional connectivity comparisons of amitriptyline (-) (control and naproxen) and amitriptyline (+) animals. Vertical bars indicate mean ± standard deviation. Two-tailed t-test or Mann-Whitney U test.

**Supplementary Figure 7 (A-B)** Remaining statistically-significant centrality alterations of each group. Note that R-FL Pagerank centrality in the control group produces a near-significant p-value (Friedman test, FDR-corrected p = 0.0567, Dunn test for multiple comparisons, p = 0.0114). Vertical bars indicate mean ± standard deviation. One-way ANOVA for repeated measures (Tukey test for multiple comparisons) or Friedman test (Dunn test for multiple comparisons). *P<0.05, **P<0.01, ***P<0.001, ****P<0.0001.

**Supplementary Figure 8 (A)** Baseline comparisons of centrality in animals with high and low CSD threshold.  **(B)** Baseline comparisons of centrality measures in amitriptyline (-) (control and naproxen) and amitriptyline (+) animals. Vertical bars indicate mean ± standard deviation. Two-tailed t-test or Mann-Whitney U test.

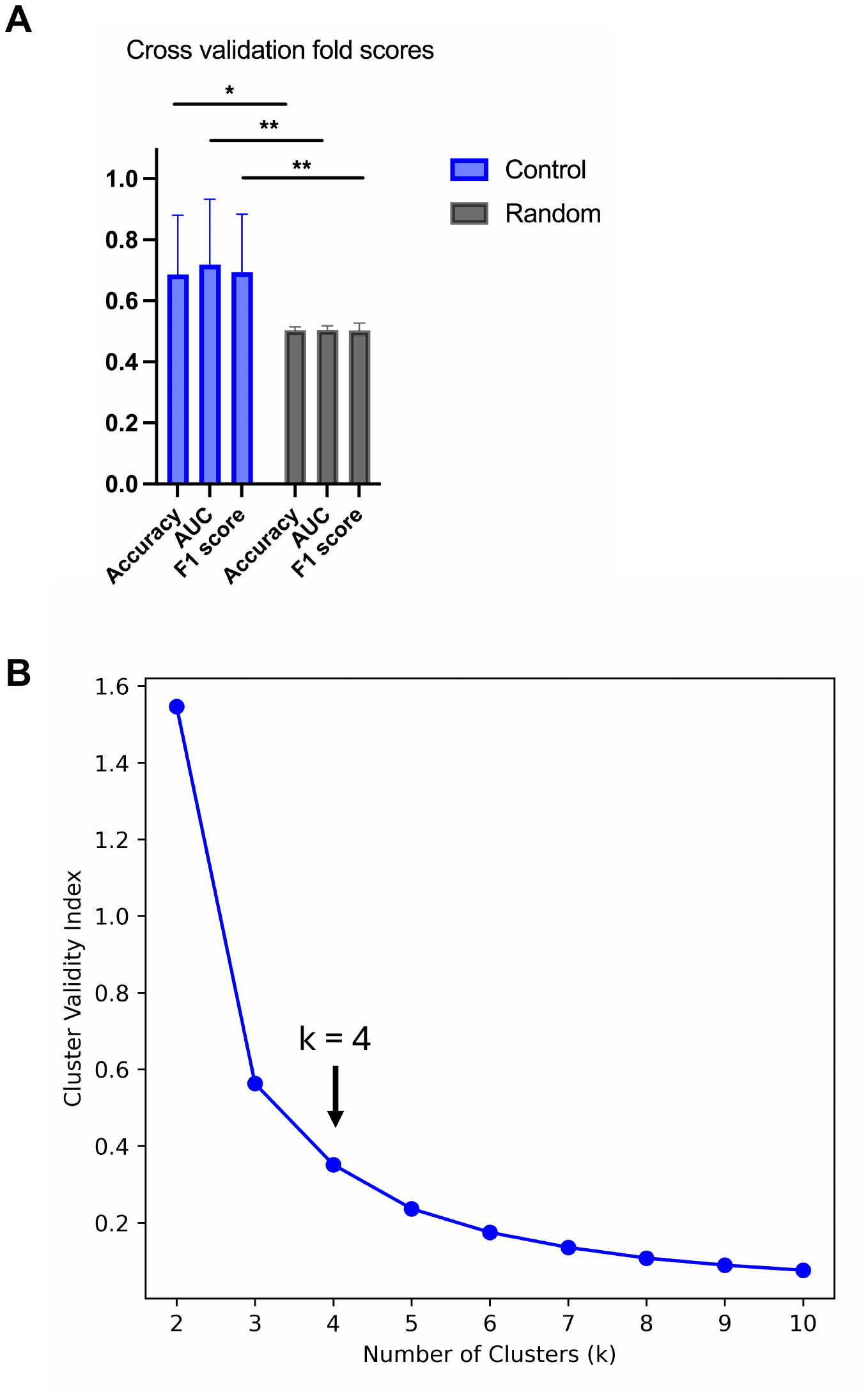

**Supplementary Figure 9 (A)** Per fold AUC, accuracy and F1 scores during cross-validation. **(B)** Elbow criterion using cluster validity index. Arrow shows the elbow when k = 4.
